# Reconstructing Spatial Transcriptomics at the Single-cell Resolution with BayesDeep

**DOI:** 10.1101/2023.12.07.570715

**Authors:** Xi Jiang, Lei Dong, Shidan Wang, Zhuoyu Wen, Mingyi Chen, Lin Xu, Guanghua Xiao, Qiwei Li

## Abstract

Spatially resolved transcriptomics (SRT) techniques have revolutionized the characterization of molecular profiles while preserving spatial and morphological context. However, most next-generation sequencing-based SRT techniques are limited to measuring gene expression in a confined array of spots, capturing only a fraction of the spatial domain. Typically, these spots encompass gene expression from a few to hundreds of cells, underscoring a critical need for more detailed, single-cell resolution SRT data to enhance our understanding of biological functions within the tissue context. Addressing this challenge, we introduce BayesDeep, a novel Bayesian hierarchical model that leverages cellular morphological data from histology images, commonly paired with SRT data, to reconstruct SRT data at the single-cell resolution. BayesDeep effectively model count data from SRT studies *via* a negative binomial regression model. This model incorporates explanatory variables such as cell types and nuclei-shape information for each cell extracted from the paired histology image. A feature selection scheme is integrated to examine the association between the morphological and molecular profiles, thereby improving the model robustness. We applied BayesDeep to two real SRT datasets, successfully demonstrating its capability to reconstruct SRT data at the single-cell resolution. This advancement not only yields new biological insights but also significantly enhances various downstream analyses, such as pseudotime and cell-cell communication.

## INTRODUCTION

Understanding the spatial distribution of transcript expression provides valuable insights into biological function and histopathology^1^. The advent of single-cell RNA sequencing (scRNA-seq) techniques has profoundly transformed our understanding of gene expression regulation in different cell lineages or types. However, tissue dissociation in scRNA-seq leads to the loss of the spatial context of gene expression, which is essential for comprehending the cell-cell and cell-environment interaction mechanism^2^. Recent advancements in spatially resolved transcriptomics (SRT) have enabled the exploration of cellular transcription in conjunction with the spatial organization and morphological features of cells. Next-generation sequencing (NGS)-based spatial molecular profiling platforms, such as Spatial Transcriptomics^3^ (ST) and 10x Visium^4^ (i.e., an improved platform by 10x Genomics), utilize spatial barcodes to capture RNA molecules, synthesize their complementary DNA molecules, and subsequently perform sequencing. These techniques facilitate the measurement of whole-genome expression across thousands of spatial locations, referred to as ‘spots’, on the tissue section. The development of SRT technology has provided valuable clinical and biological insights into various areas, including tumor heterogeneity, brain function, and sepsis pathophysiology^5-7^.

One of the primary constraints inherent to NGS-based SRT techniques is their spatial resolution. Gene expression in these platforms is measured on an array of spots, with a typical diameter of 100 μ*m* for ST and 55 μ*m* for 10x Visium^8^. Consequently, depending on the platform and the type of biological tissue being analyzed, each spot’s area may encompass a heterogeneous population of cells, ranging from a few to as many as 200 cells. Furthermore, SRT techniques solely measure gene expression within the area of the confined array spots, thus covering part of the total cells observed within the entire tissue section. To illustrate, the area covered by measured spots accounts for approximately 1/3 of the total tissue region for the 10x Visium platform and around 20% for ST platforms (see Supplementary Figure S1 for a detailed explanation). The inherent limitation in spatial resolution and the scope of regions could lead to substantial information loss when analyzing SRT data. Such limitations may, in turn, impact the applicability of these techniques in facilitating the discovery of deeper insights in various biomedical studies, specifically in investigating gene expression patterns at the cellular level.

Several computational methods have been developed to enhance the spatial resolution of SRT data, addressing these limitations. For instance, BayesSpace^8^ employs a Bayesian approach to improve gene expression at the sub-spot resolution. However, despite this advancement, sub-spots generated by BayesSpace may still encompass multiple cells, yielding only a modest improvement in spatial resolution. Furthermore, BayesSpace cannot predict gene expression outside the spotted regions. In parallel, a series of approaches, such as TESLA^9^, XFuse^10^, ST-Net^11^, and HisToGene^12^, have been developed to reconstruct gene expression at super-resolution. These deep-learning-based techniques utilize molecular information from SRT data and morphological information from the paired histology image. Notably, while these methods enhance gene expression prediction at the pixel level, which may complicate interpretation, none directly account for cellular spatial organization in their analyses.

In response, we introduce a novel Bayesian methodology, BayesDeep, for deeply resolving gene expression for all “real” cells by integrating the molecular profile from SRT data and the morphological information extracted from the paired histology image. Specifically, BayesDeep builds upon a regularized negative binomial regression model with grouped observations. The response variable is the spot-resolution gene expression measurements in terms of counts; and the explanatory variables are a range of cellular features extracted from the paired histology image, including cell type and nuclei-shape descriptors. Following the estimation of regression coefficients, BayesDeep predicts the gene expression of all cells based on their cellular features, regardless of whether they are within or beyond spot regions. The model robustness is achieved by regularization using a spike-and-slab prior distribution to each regression coefficient. We validated the accuracy of gene expression prediction at both spot and single-cell resolution using simulated and real SRT datasets. Additionally, we demonstrated that the single-cell-resolution spatial molecular profiles characterized by BayesDeep enable in-depth investigations on cell-type specific differential expression analysis, cell-cell communication analysis, and pseudo-time analysis. Overall, our findings highlight the effectiveness of BayesDeep in reconstructing molecular profiles at the single-cell resolution from NGS-based SRT data, which is typically limited to spot resolution, paving the way for advanced research in pseudotime and cell-cell communication analysis.

## RESULTS

### Overview of BayesDeep

BayesDeep builds upon a Bayesian negative binomial regression model to recover gene expression at the single-cell resolution from NGS-based SRT data. The schematic diagram is depicted in Figure 1.

**Figure 1.**
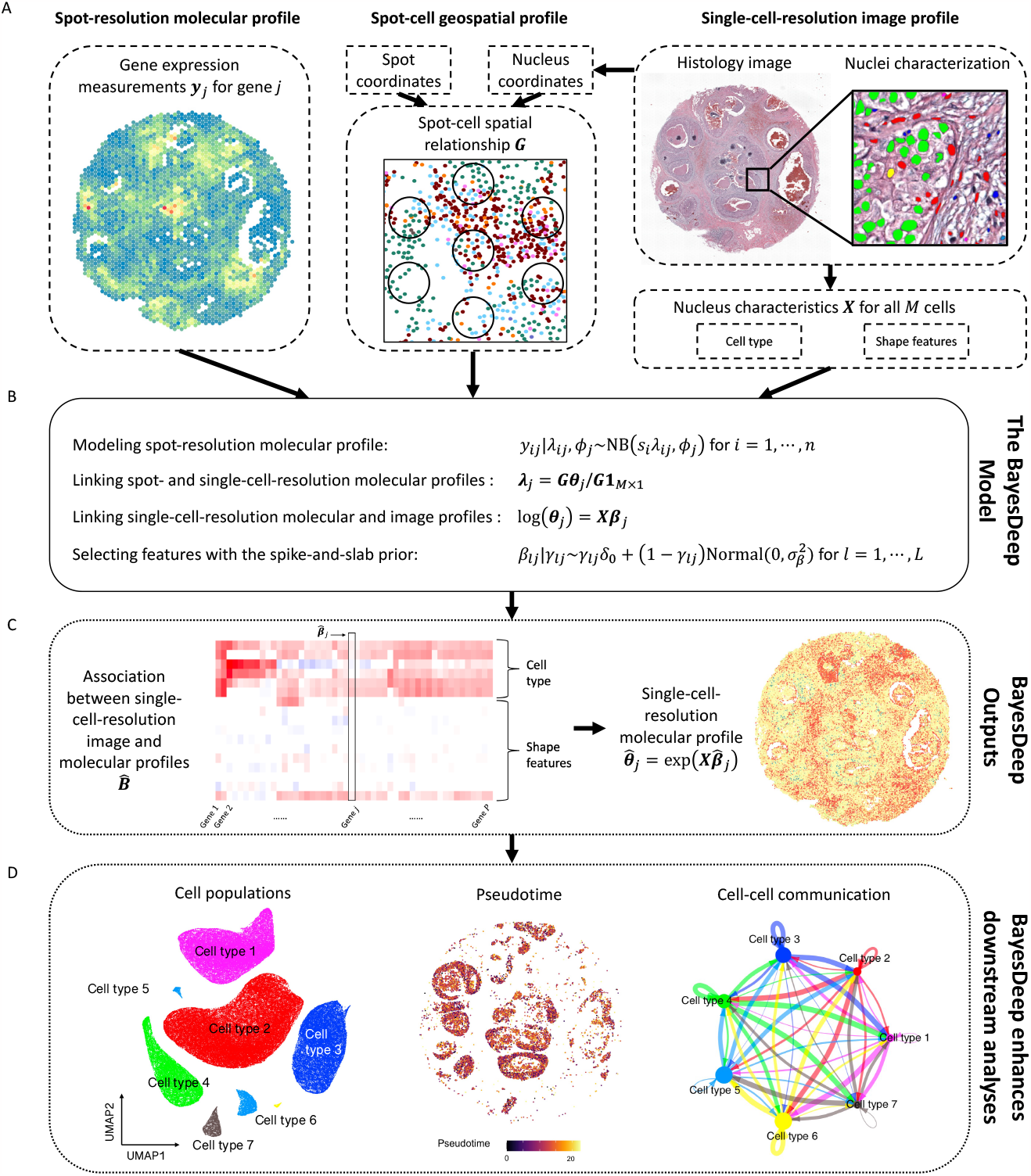
Flowchart of the proposed BayesDeep: A. BayesDeep integrates the spot-resolution molecular profile ***Y*** from NGS-based SRT data, the single-cell-resolution image profile ***X*** from the paired AI-reconstructed histology image, and the spot-cell geospatial profile ***G*** to recover gene expression at the single-cell resolution ***Θ***. B. The hierarchical formulation of the BayesDeep model is based on a Bayesian regularized negative binomial regression model with grouped observations. C. BayesDeep estimates the association between the single-cell-resolution molecular and image profiles ***B*** and predicts the single-cell-resolution molecular profile ***Θ***. D. Several downstream analyses can be enhanced based on the availability of the single-cell-resolution molecular profile ***Θ***, including identifying distinct cell populations, elucidating the process of tumorigenesis *via* pseudotime analysis, and exploring the mechanisms of cell-cell communication.

BayesDeep integrates three distinct modalities from a standard NGS-based SRT experiment: the molecular, image, and geospatial profiles (see Figure 1A). The molecular profile refers to the spot-resolution gene expression data denoted by an *N*-by-*P* count matrix ***Y***, where *N* is the number of spots and *P* is the number of genes. The image profile corresponds to the detailed morphological context of the paired histology image in terms of a set of cellular features. We use any *M*-by-*L* design matrix ***X*** to denote the image profile, where *M* is the number of all observed cells and *L* is the number of cellular features, which may include cell types, nuclei-shape characteristics, and any other relevant explanatory features that can be quantified at large scale. The geospatial profile reveals the spatial relationship between the *N* spots and *M* cells, which can be defined by an *N*-by-*M* binary matrix ***G***, with one signifying that a cell is within the barcoded area of a spot.

The spot-resolution gene expression matrix ***Y*** and the single-cell-resolution morphological features of those cells within spot regions serve as a reference for recovering the single-cell-resolution gene expression of all *M* cells, whether within or beyond spot regions. The model is specified in Figure 1B. We first modeled the observed read count for a specific gene within a spot using a negative binomial (NB) distribution. Then, the underlying spot-resolution relative gene expression in the NB mean is assumed to be the average of single-cell-resolution relative expression across all cells within the spot. Next, we considered the logarithm of each cell’s relative expression as a linear combination of covariates that includes a scalar of one for the intercept and *L* measurable explanatory variables that pertain to that cell. A spike-and-slab prior model is applied for each covariate coefficient. On one hand, this feature selection scheme improves the robustness of our model. On the other hand, the corresponding coefficient matrix ***B*** uncovers significant associations between gene expression and cellular characteristics, illustrated in Figure 1C, thereby potentially offering valuable biological insights. With the reconstructed single-cell-resolution spatial molecular profile ***Θ***, we can undertake several pivotal downstream analyses, as depicted in Figure 1D. These analyses allow for the differentiation of cell populations, the exploration of tumorigenesis through pseudotime analysis, and the dissection of ligand-receptor signaling pathways vital for cell-cell communication.

### Model Validation

#### Model validation on simulated data

We designed a simulation study to validate the accuracy of BayesDeep. We selected a connected region of *N* = 500 spots from the human breast cancer 10x Visium data (displayed as green circles in Figure 2A) to generate the spot-resolution molecular profile. The single-cell-resolution image profile ***X***, including cell types and nuclei-shape features and the locations of cells, were from the nuclei identification results by HD-staining^13^ for the paired histology image of the SRT data. As introduced in the Method section and observed in real SRT data analysis, the covariate coefficient matrix ***B*** is highly sparse, indicating that many explanatory variables do not contribute to gene expression for most genes. To replicate this condition, we generated the coefficients, ensuring nearly half were zeros, reflecting the observed zero-inflation. Based on the model assumption of BayesDeep, we first generated single-cell-resolution relative gene expression matrix ***Θ*** on the selected region, and then the spot-resolution gene expression matrix ***Y*** on all spots within. The data-generating procedure is detailed in section S1 in Supplementary Materials.

**Figure 2.**
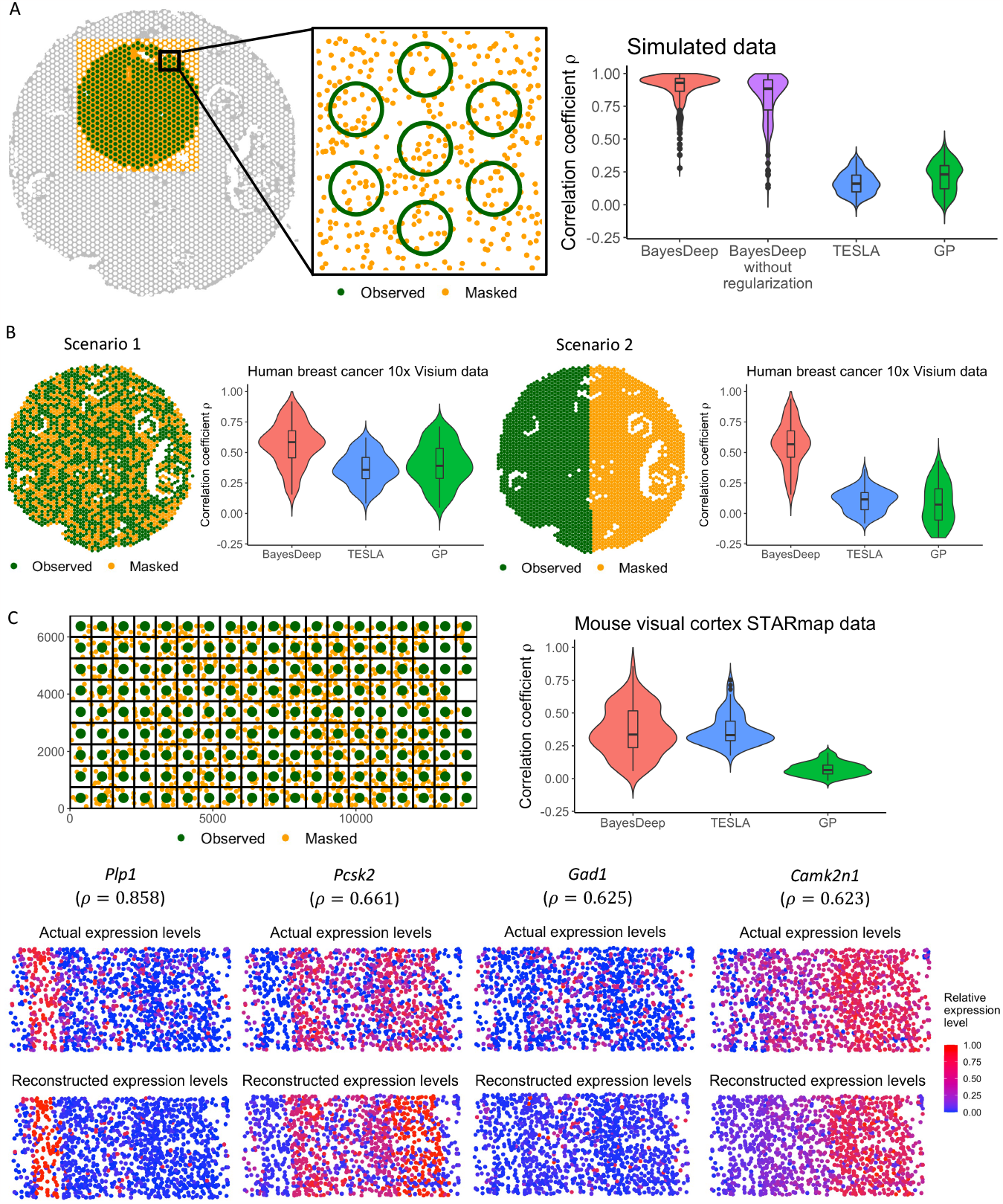
Overview of model validation, including the validation settings and evaluation results in terms of the Pearson correlation coefficients ρ between the actual and predicted gene expression for BayesDeep, TESLA, and Gaussian Process (GP), respectively. The validation is stratified into three distinct data: A. Simulated data at the spot resolution; B. Human breast cancer 10x Visium data at the spot resolution; C. Mouse visual cortex STARmap data at the single-cell resolution.

We evaluated the performance of BayesDeep in recovering the single-cell-resolution molecular profile ***Θ***, against two other methods, TESLA^9^ and Gaussian process (GP)^14^, by measuring the Pearson correlation coefficient *ρ* ∈ [−1,1] between the logged estimated and actual gene expression at the single-cell resolution, 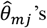 and *θ*_*mj*_’s. Using relative gene expression allows us to discern which cells exhibit high gene expression and which demonstrate lower expression. A high positive *ρ* value signifies the successful model outcome and robust overall performance. As depicted in Figure 2A, the results demonstrate that BayesDeep, when employing regularization, significantly surpasses the comparative methods, with a median ρ of 0.928. In contrast, TESLA and GP failed to reconstruct the single-cell-resolution molecular profile ***Θ***, as indicated by their respective ρ values approaching zero. To further examine the efficacy of the regularization approach, we computed the root mean square error (RMSE) between the estimated and actual covariate coefficients, 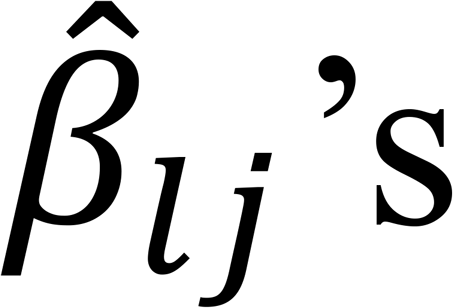 and *β*_*lj*_’s, with lower RMSE values denoting more precise estimations. Specifically, in scenarios where regularization was omitted, a standard normal prior was applied in place of the spike-and-slab prior to each covariate coefficient *β*_*lj*_. These results demonstrate that the incorporation of regularization significantly refines the precision of estimations for non-zero coefficients and substantially improves accuracy for truly zero coefficients (Supplementary Figure S2). These findings support the implementation of a regularized prior within our BayesDeep framework.

#### Model validation on real SRT data at the spot resolution

In this study, we utilized the human breast cancer^15^ and human prostate cancer 10x Visium data^16^ in the real data analysis to demonstrate the efficacy of BayesDeep in restoring gene expression within manually masked spots. For each SRT data, we randomly selected *P* = 100 genes with non-zero read counts in at least 50% spots. We assessed BayesDeep against TESLA and GP, through two distinct masking scenarios for spot-resolution gene expression imputation. In scenario 1, 40% of spots were randomly masked, while in scenario 2, we divided the domain into two halves and masked all spots on one side (refer to Figure 2B, with masked spots marked in orange and unmasked spots marked in green). The second scenario aimed to challenge the model’s predictive power over a separate region that does not overlap with those masked spots. The gene expression of those remaining unmasked spots ***Y*** served as the main input of each method. For BayesDeep, we also incorporated the morphological information ***X***, detailed in the Method section, from the corresponding histology image.

Figure 2B shows the correlation coefficient ρ for comparing observed and imputed spot-resolution gene expression in the masked spots within the human breast cancer 10x Visium data. BayesDeep achieved a median correlation of 0.585 in scenario 1, significantly higher than that of the competing methods (TESLA: ρ = 0.358; GP: ρ = 0.391). In scenario 2, the performance of BayesDeep is comparable to scenario 1, whereas TESLA and GP failed to impute gene expression for the entirely masked half, as evidenced by negligible correlation values. This failure is attributed to their spatial dependence assumption in reconstructing gene expression, resulting in inadequate performance when the masked and observed regions do not intersect. Supplementary Figure S3 shows similar results through the analysis on human prostate cancer 10x Visium data. These results show the capability of BayesDeep to predict gene expression at the spot resolution, both in the same and new regions, by leveraging single-cell-resolution morphological features.

#### Model validation on real SRT data at the single-cell resolution

We extended our study to assess the performance of BayesDeep on the prediction of single-cell-resolution gene expression when ground truth is available at the single-cell resolution. We validated the model using the mouse visual cortex STARmap data^17^, which was collected by an imaging-based SRT platform. It comprises measurements of 1,020 genes across *M* = 1,207 cells, categorized into *Q* = 15 distinct cell types distributed across seven layers labeled in the original study. In our analysis, we filtered out genes that are expressed in less than 30% cells or the highest read count is less than ten. With these filtering criteria, we kept *P* = 77 genes for the following analysis. We then constructed the spot-resolution molecular profile ***Y*** by overlaying a square lattice grid across the entire domain, with each square unit representing a “spot” of 750 × 750 pixels, as illustrated in Figure 2C. This yielded *N* = 105 spots, each containing more than one cell, covering the entire domain without inter-spot gaps. The read count *y*_*ij*_ of gene *j* for spot *i* was calculated by aggregating the read counts from all cells within that spot. Owing to the absence of a corresponding histology image for the data, we utilized the cell type and layer information provided along with the SRT data in the original study as explanatory variables ***X*** to inform BayesDeep.

Figure 2C visualizes the correlation coefficient ρ between the imputed and true relative expression across all cells for all genes. BayesDeep achieved a median correlation of approximately 0.336, similar to TESLA (median ρ = 0.331) but significantly outperforming GP (median ρ = 0.065). Additionally, Figure 2C illustrates the actual *versus* predicted relative expression for four representative genes, demonstrating a notable agreement between the actual and predicted gene expression patterns. These findings affirm the capability of BayesDeep to reconstruct the expression patterns of the underlying cells from spot-resolution gene expression accurately.

### Application to Human Breast Cancer 10x Visium Data

We applied BayesDeep to reconstruct the single-cell-resolution molecular profiles for the SRT data from a human breast cancer study^15^. The data includes *N* = 2,518 spots and 17,651 genes. The gene expression was measured on a section of the human breast with invasive ductal carcinoma *via* the 10x Visium platform, along with annotation from pathologists as a reference (H&E-stained image with five annotated tissue regions in Figure 3A). After applying HD-Staining^13^ to the histology image of breast cancer tissue, we identified *M* = 156,235 cells within seven categories: macrophage, ductal epithelium, karyorrhexis, tumor cell, lymphocyte, red blood cell, and stromal cell. Supplementary Table S1 provided ten shape features used in this study. We applied BayesDeep to reconstruct the single-cell-resolution molecular profiles on the top *P* = 2,000 highly variable genes, compared its performance to a competing method, TESLA, and further utilized the generated higher-resolution molecular profiles on several downstream analyses to reveal biological insights.

**Figure 3.**
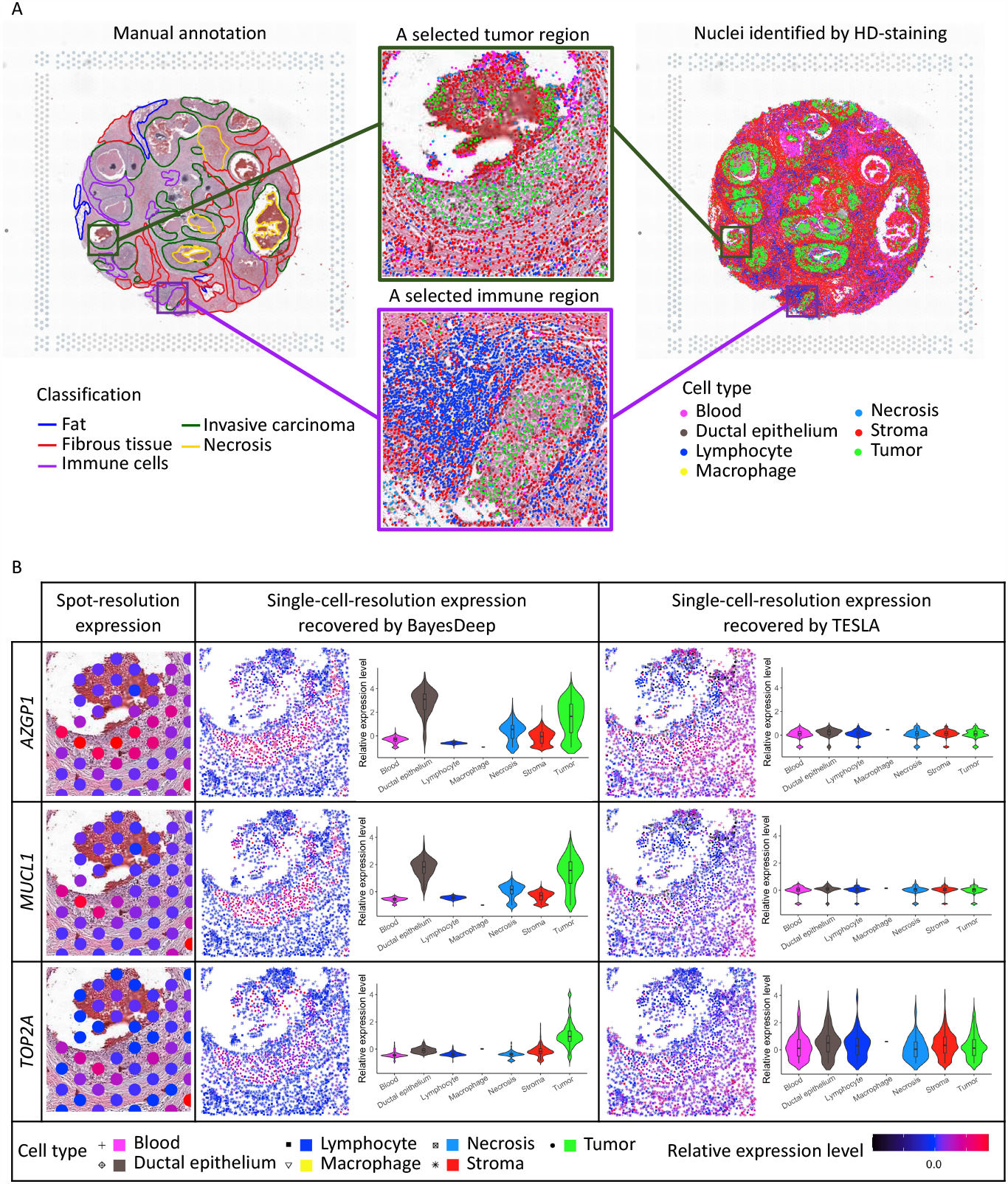
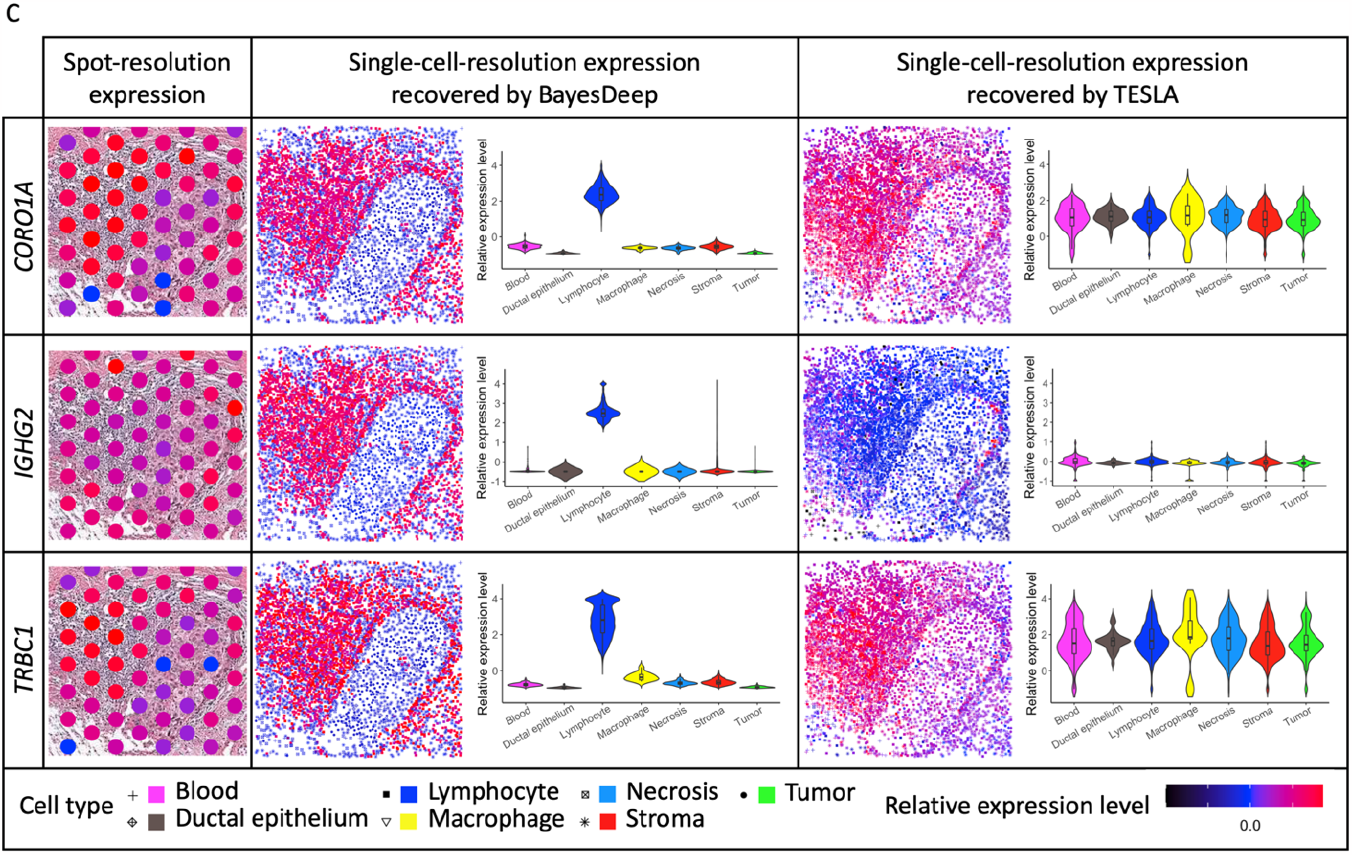
Real data analysis on the human breast cancer 10x Visium data: A. Manual annotation and nuclei identification results by applying HD-staining to the paired histology image; and the two selected tumor and immune regions for an illustrative comparison between BayesDeep and TESLA. B. The actual expression at the spot resolution and the predicted expression at the single-cell resolution by BayesDeep and TESLA on the selected tumor region for genes *AZGP1, MUCL1*, and *TOP2A*. Violin plots display gene expression on the selected tumor region across different cell types. C. The actual expression at the spot resolution and the predicted expression at the single-cell resolution by BayesDeep and TESLA on the selected immune region for genes *CORO1A, IGHG2*, and *TRBC1*. Violin plots display gene expression on the selected tumor region across different cell types.

The gene expression at the single-cell resolution generated by BayesDeep offers a more detailed view of the spatial transcriptome landscape within the cellular environment. We chose both a tumor and an immune-related region as illustrative examples to demonstrate BayesDeep’s capacity to enhance our understanding of gene expression in regions with high cell type heterogeneity (Figure 3A). For the example tumor region, we observed two distinct small areas densely populated with tumor cells. We further examined and presented the original spot-resolution expression from SRT data and reconstructed expression using BayesDeep and TESLA of three breast cancer-related genes (Figure 3B): First of all, *AZGP1*, which is primarily expressed in breast epithelial cells, plays a multifaceted role associated with cancer cachexia, carcinogenesis, and tumor differentiation^18^. Secondly, *MUCL1*, a breast-specific gene predominantly expressed in breast cancer, serves as a vital biomarker for tumor progression and metastasis^19,20^. Last but not least, *TOP2A* serves as a notable proliferation marker, demonstrating high expression in various subtypes of breast cancers^21^. The detailed expression pattern is relatively difficult to observe at the spot resolution (spot-resolution relative expression levels in Figure 3B) due to the low resolution. However, the expression reconstructed by BayesDeep for these three example genes distinctly exhibits the expression pattern consistent with the distribution of tumor cells, i.e., high expression on tumor cells and low expression on other cells. Conversely, for the gene expression reconstructed by TESLA, no similar expression pattern is observed in the two tumor areas for all three genes, indicating its limitation in recovering the comprehensive cellular expression. Violin plots in Figure 3B show the higher expression on tumor cells and ductal epithelium for BayesDeep, which aligns with the gene functions and their association with breast cancer. In contrast, the predicted expression from TESLA shows no difference among different cell types. Another observed region in Figure 3A is an immune region predominantly comprised of lymphocytes, which covers around 56 spots in ST data. In this region, many lymphocytes encircle a small tumor area. We selected three example genes related to immune functions for validation purposes. *CORO1A* is an identified immunity gene signature in breast cancer cohort studies^22^. *IGHG2* encodes the constant region of the immunoglobulin gamma-2 heavy chain, which enables antigen binding activity and immunoglobulin receptor binding activity and is involved in several processes, such as activating immune response^23^. *TRBC1* encodes T cell receptor *β* chain constant region 1, which is partially expressed in subsets of T cells^24,25^. No obvious expression patterns can be observed at the spot resolution due to the limited resolution (Figure 3C). However, through single-cell-resolution reconstruction *via* BayesDeep, we observed that these three genes exhibited high expression in the surrounding immune area. Again, gene expression recovered by TESLA shows no difference among various cell types and spatial regions. In summary, BayesDeep-predicted single-cell-resolution gene expression exhibits significantly stronger associations with the cell type information, unveiling detailed gene expression patterns not captured by the ST data alone.

In addition, BayesDeep quantifies the association between gene expression and the cellular features extracted from the histology image by estimating the covariate coefficient matrix ***B***. In Figure 4A, the heatmap displays the coefficients of seven distinct cell types and ten nuclei-shape descriptors for the top 2,000 highly variable genes. It is noticed that a substantial number of the estimated covariate coefficients *β*_*lj*_’s are non-zero and vary across cell types for most genes, which highlights unique expression patterns for different cell types. On the other hand, certain genes are characterized by all *β*_*lj*_ ‘s being zero, implying homogenous expression across cell types. In contrast, the covariate coefficients of those shape descriptors excluding solidity are notably sparse, with approximately 87.0% of those coefficients being zero. There exists an association between gene expression and the solidity of the cell nucleus for around 50.1% of genes. It is important to note that the orientation covariate (i.e., the third column from the right of the heatmap shown in Figure 4A) is a negative control. This covariate is expected to be independent of gene expression, given that the measurement of nuclear orientation relies on the tissue’s placement on the slide. We observed that the estimated coefficients for the orientation covariate are exactly zero for all genes, indicating the validity of BayesDeep results.

**Figure 4.**
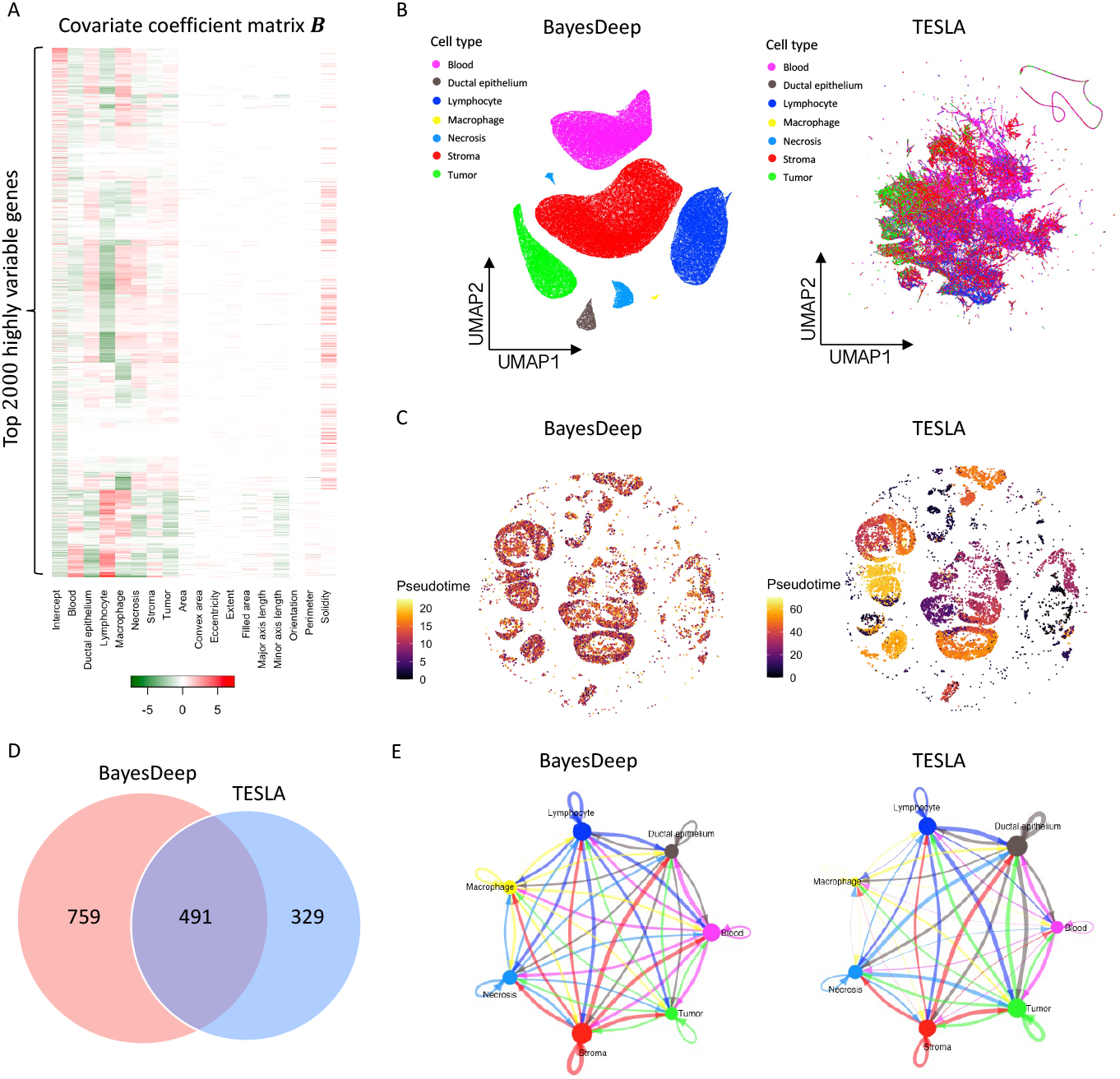
Real data analysis (continued) and downstream analysis on the human breast cancer 10x Visium data: A. Heatmap of the covariate coefficient matrix ***B*** estimated by BayesDeep, indicating the association between gene expression and morphological features extracted from the paired histology image. B. Cell population analysis on the BayesDeep- and TESLA-generated single-cell-resolution gene expression. C. Pseudotime analysis on tumor cells extracted from the cell population analysis. D and E. Cell-cell interactions inferred from the BayesDeep- and TESLA-generated single-cell-resolution gene expression, and their overlap.

To demonstrate that BayesDeep-reconstructed single-cell-resolution gene expression data can improve downstream analysis, we conducted a comparative analysis between BayesDeep and TESLA across three tasks: (i) distinguishing different cell populations, (ii) explicating tumorigenesis through pseudotime analysis and (iii) elucidating cell-cell communication patterns. First, we employed Seurat for clustering and dimensional reduction analysis on the BayesDeep-generated single-cell-resolution gene expression. The results demonstrated that cells of the same cell type co-localized well on the UMAP plot and exhibited clear separation among distinct cell types (Figure 4B left panel). In contrast, we performed a similar analysis on the gene expression generated by TESLA. However, the results showed that TESLA could not effectively separate many cell types (Figure 4B right panel). These findings underscore the capability of BayesDeep to distinguish different cell populations. Second, we employed Moncle3^26,27^ to perform a pseudotime analysis on tumor cells extracted from BayesDeep and TESLA results. In such analysis, we incorporated *CD44* and *CD24* as breast cancer stem cell markers, as previously reported^28-30^. We designated the cells with the highest *CD44* and *CD24* expression as the root of the trajectory to assess cancer progression over pseudotime (Supplementary Figure S4). Predicted results based on BayesDeep indicated that cancer cells from earlier pseudotime are predominantly enriched in the conventional tumor region. In contrast, those from later pseudotime are primarily concentrated within the necrotic tumor regions^31^ (Fig. 4C, left panel). This trajectory pattern is consistent with the timing order of tumor evolution. In contrast, the pseudotime derived from TESLA-generated predictive results did not show a biologically meaningful trend (Figure 4C, right panel). Overall, our results demonstrate the consistent superiority of BayesDeep in explicating tumorigenesis through the pseudotime analysis. Third, because BayesDeep can identify various cell types more accurately than TESLA, we hypothesized that BayesDeep could offer more details of cell-cell communications among these cell types than TESLA. To corroborate this hypothesis, we applied CellChat^32^ to define the cell-cell communication landscape based on the cell clusters identified by pathologist annotations. Both BayesDeep and TESLA can achieve abundant cell-cell interactions, but BayesDeep can reveal more cellular communications than TESLA (Figure 4D). Interestingly, the majority (1,001/1,486 = 67.3%) of cell-cell communications revealed by BayesDeep overlapped with those identified by TESLA (Figure 4D). However, some detailed cell-cell interactions revealed by BayesDeep and TESLA are different (Figure 4E): for example, BayesDeep could identify more immune-related, especially macrophage-related cellular communications than TESLA. In general, consistent with our expectations, BayesDeep can reveal more cell-cell communications than TESLA.

### Application to Human Prostate Cancer 10x Visium Data

To assess the adaptability of BayesDeep across diverse tissue types, we conducted analysis utilizing another SRT data derived from human prostate cancer tissue^16^. The data includes *N* = 4,371 distinct spots for 17,651 genes. The gene expression measurement was obtained from a section of invasive carcinoma within the human prostate (H&E-stained image with six annotated tissue regions in Figure 4A), utilizing the 10x Visium platform. Subsequently, we applied the HD-Staining technique to identify nuclei on the histology image of this tissue. This image analysis process led to the segmentation of *M* = 259,257 individual cells, which were systematically categorized into six classes: macrophage, karyorrhexis, tumor cell, lymphocyte, red blood cell, and stromal cell. We utilized BayesDeep to reconstruct high-resolution molecular profiles for the top 2,000 highly variable genes at the single-cell resolution. We then assessed its performance against the TESLA method and leveraged the resulting detailed molecular profiles in several downstream analyses.

To examine the gene expression reconstructed at the single-cell resolution by BayesDeep and TESLA, we selected two example regions with a high cell-type mixture - a tumor and an immune-related region (in Figure 5A). For the example tumor region depicted in Figure 5A, we observe that tumor cells have circular patterns around each empty region. We examined the expression of three prostate cancer-related genes (Figure 5B). *ADGRF1* is an adhesion-G protein-coupled receptor and has an essential function in cancer^33^. Aberrant expression and mutation of G protein-coupled receptors and their signaling partners, G proteins, have been well documented in many cancer^34^. *SPON2* (tumor cell-derived spondin 2) is an extracellular matrix glycoprotein, and overexpression of *SPON2* has been shown to promote tumor cell migration^35^. *TMEFF2* encodes a transmembrane protein containing an epidermal growth factor-like motif and two follistatin domains, highly expressed in prostate cancer samples ^36^. At the spot resolution in Figure 5B, the expression patterns of these genes remain obscure. However, the gene expression reconstructed by BayesDeep shows strong expression patterns. These three example genes have higher expression in tumor cells. In contrast, the gene expression reconstructed by TESLA does not display differential expression on tumor cells. The violin plots in Figure 5B provide evidence of the differential expression among cell types for BayesDeep, in line with the three example genes’ cancer-related functions. For another selected region, as depicted in Figure 5A, we find an immune region in the center primarily composed of lymphocytes. For our analysis, we selected three immune-related genes. *CD24* is a cell surface glycosyl-phosphatidylinositol–anchored protein expressed on various cell types, including developing T and most B lymphocytes^37^. *CD47* is an immunoglobulin superfamily pentatransmembrane protein ubiquitously expressed in hematopoietic cells, including lymphocytes^38^. *CXCR4* is the receptor for the CXC chemokine stromal-derived-factor-1, which has essential functions on immunological functions and T lymphocyte trafficking^39^. At the spot resolution, differential expression in the small immune region is not readily observable since the region is only covered by around five spots. However, through single-cell-resolution reconstruction with BayesDeep, we can clearly see that these three genes exhibit high expression within the immune area. And violin plots show their high expression within immune cells, such as lymphocytes or macrophage. In contrast, TESLA fails to detect gene expression changes that occur only in a small region.

**Figure 5.**
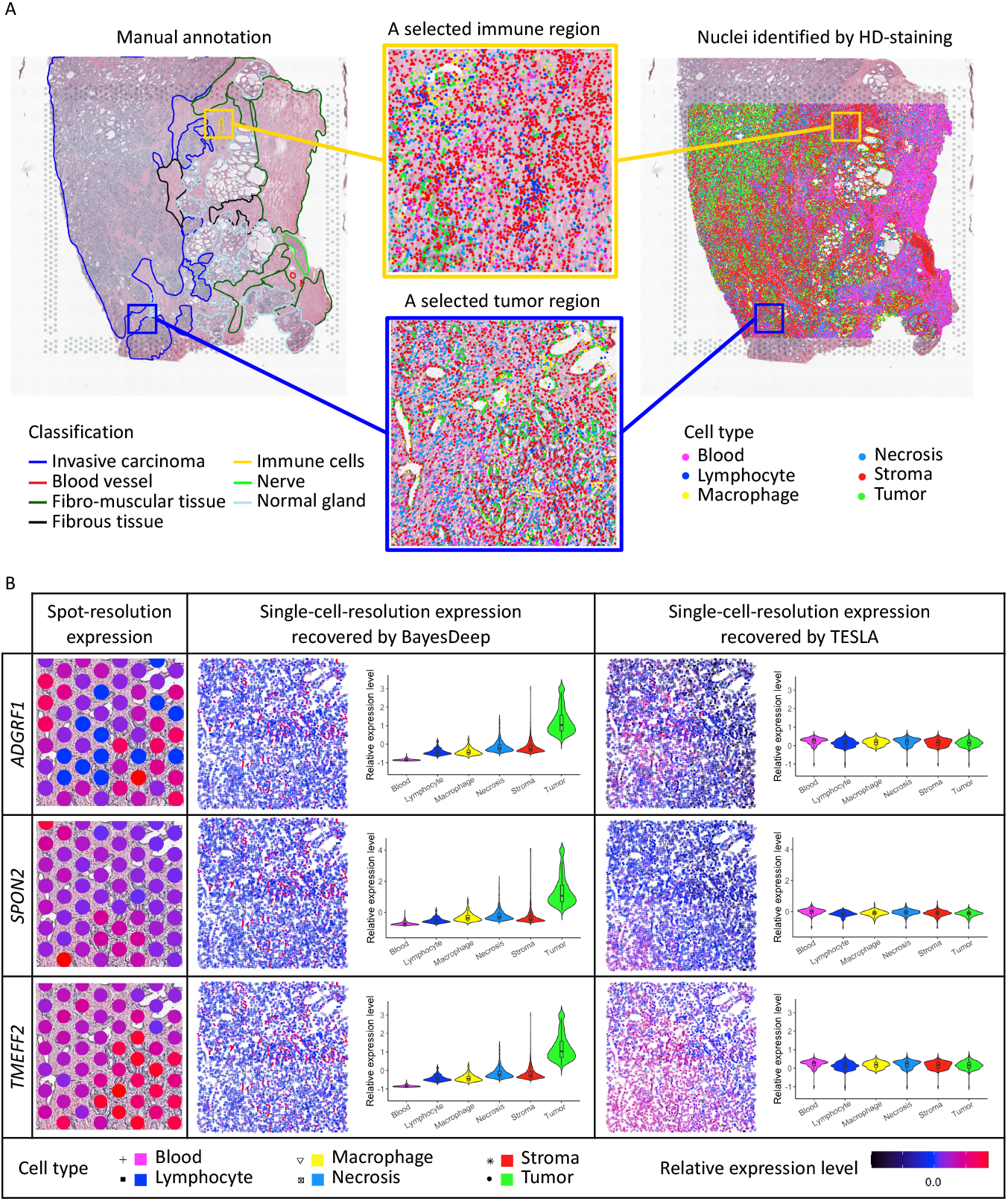
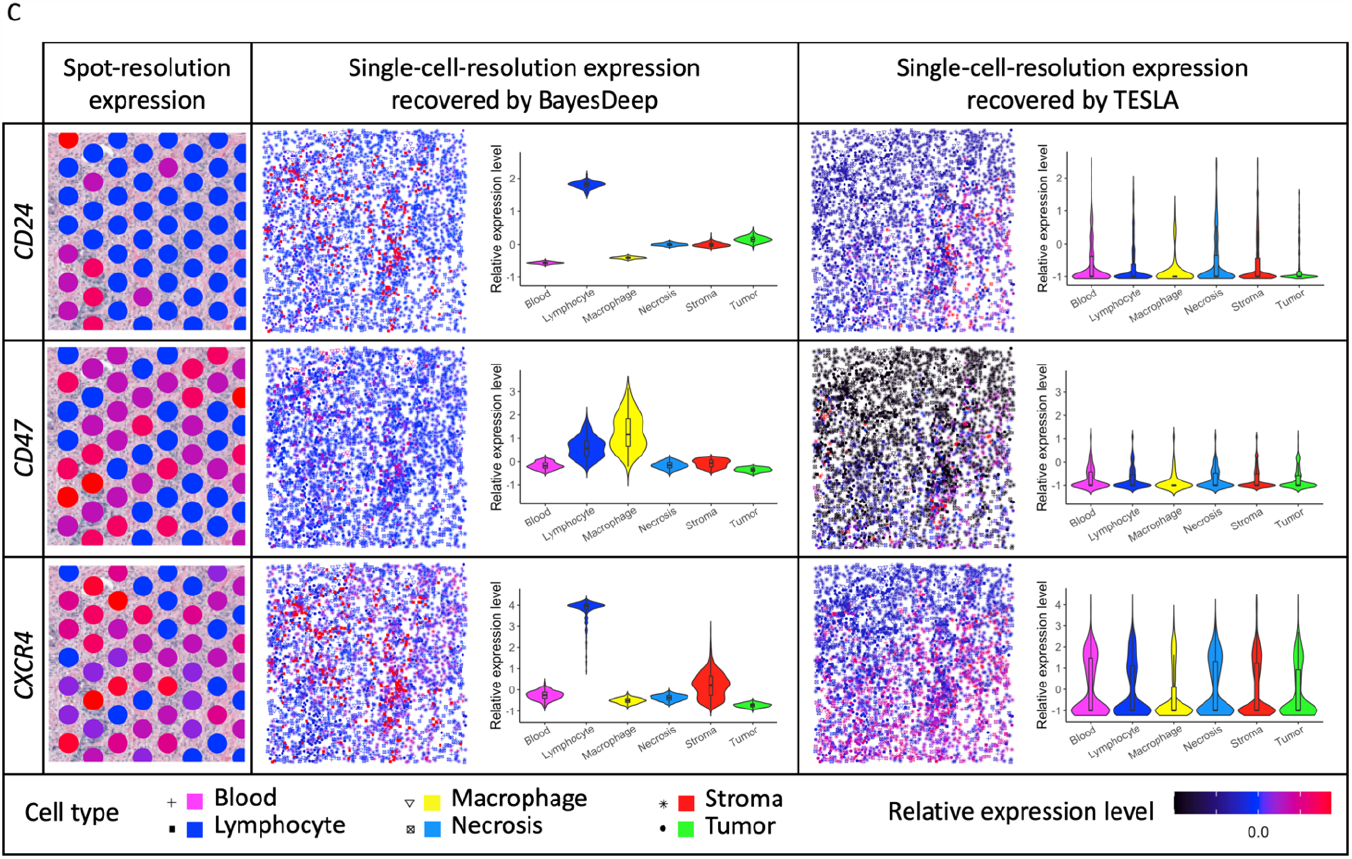
Real data analysis on the human prostate cancer 10x Visium data: A. Manual annotation and nuclei identification results by applying HD-staining to the paired histology image and the two selected tumor and immune regions for an illustrative comparison between BayesDeep and TESLA. B. The actual expression at the spot resolution and the predicted expression at the single-cell resolution by BayesDeep and TESLA on the selected tumor region for genes *ADGRF1, SPON2*, and *TMEFF2*. Violin plots display gene expression on the selected tumor region across different cell types. C. The actual expression at the spot resolution and the predicted expression at the single-cell resolution by BayesDeep and TESLA on the selected immune region for genes *CD24, CD47*, and *CXCR4*. Violin plots display gene expression on the selected tumor region across different cell types.

The inference of covariate coefficients in BayesDeep reflects the correlation between single-cell-resolution molecular profiles and extracted cell characteristics from histology images. Figure 6A shows a heatmap of coefficients for six cell types and ten nuclei-shape covariates among the top 2,000 highly variable genes. Most genes exhibit non-zero coefficients for at least one cell type, indicating the differential expression of these cell types. For nuclei-shape features, the estimated coefficients are highly sparse, with around 90.4% being zero. The estimated coefficients on the covariate for negative control and orientation are exactly zero for all genes, further validating the efficiency of BayesDeep.

**Figure 6.**
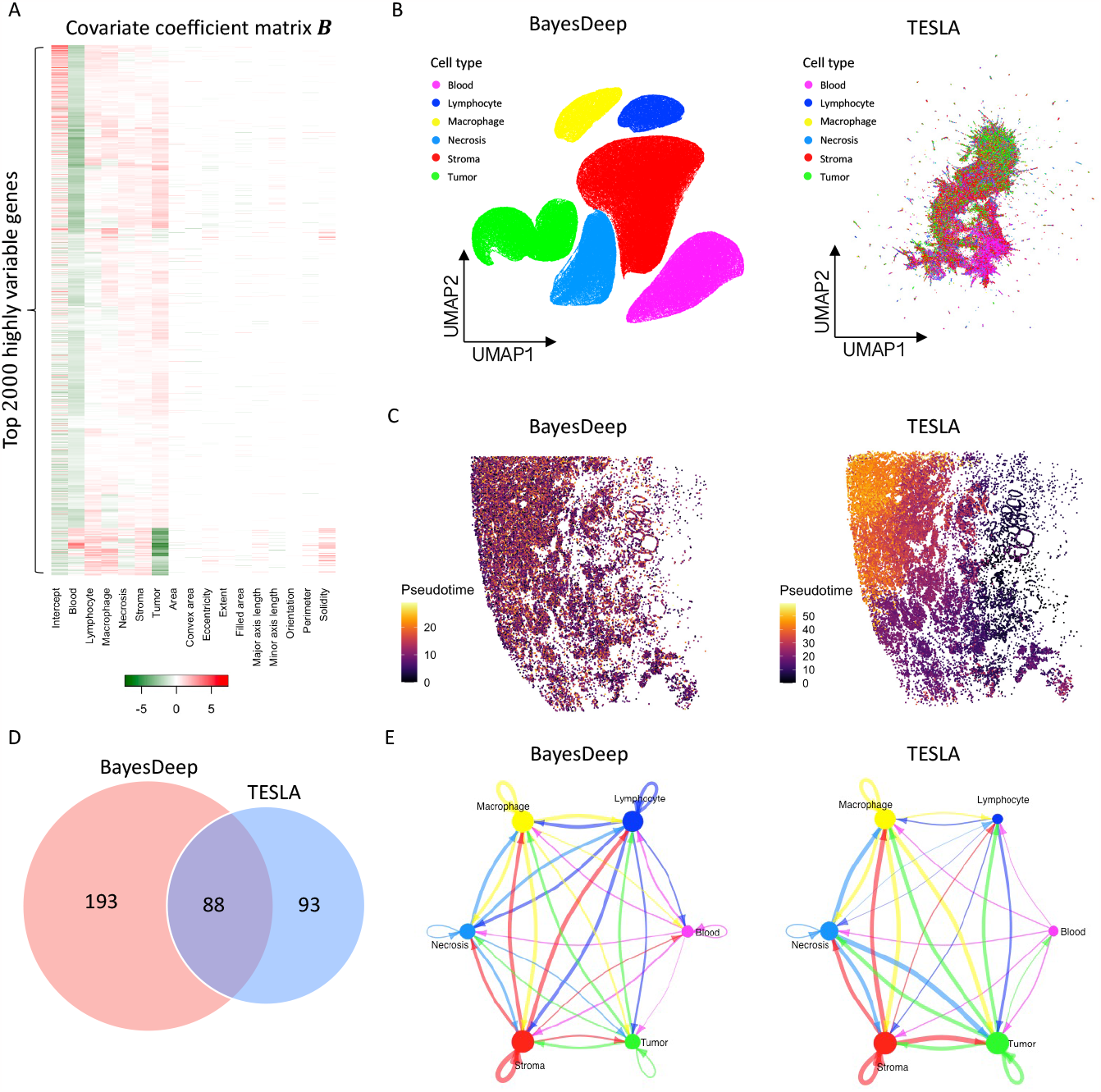
Real data analysis (continued) and downstream analysis on the human prostate cancer 10x Visium data: A. Heatmap of the covariate coefficient matrix ***B*** estimated by BayesDeep, indicating the association between gene expression and morphological features extracted from the paired histology image. B. Cell population analysis on the BayesDeep- and TESLA-generated single-cell-resolution gene expression. C. Pseudotime analysis on tumor cells extracted from the cell population analysis. D and E. Cell-cell interactions inferred from the BayesDeep- and TESLA-generated single-cell-resolution gene expression and their overlap.

Furthermore, we compared BayesDeep and TESLA on their impact on downstream analysis. First, through clustering and dimensional reduction analysis of the prostate cancer single-cell-resolution results generated by BayesDeep and TESLA, we found that the BayesDeep UMAP plot, but not the TESLA UMAP plot, revealed clear clustering results for various cell types (Figure 6B). Second, in the pseudotime analysis of prostate cancer, we used *ITGA6* and *ALCAM* as prostate cancer stem cell markers, as previously reported^28-30^ (Supplementary Figure S5). Trajectory analysis of the prostate cancer cells, extracted from BayesDeep results, revealed an even distribution of cancer cells with varying pseudotime throughout the prostate tumor. In contrast, TESLA results indicated a preference for prostate cancer cells with earlier pseudotime to be enriched at the tumor periphery (Figure 6C). Third, we assessed BayesDeep’s ability to identify cell-cell communication in prostate cancer tissue compared to TESLA. BayesDeep identified 281 ligand-receptor interacting pairs, more than twice the number identified by TESLA (Figure 6D). In summary, these findings demonstrated that BayesDeep could outperform TESLA in multiple downstream analysis tasks and, therefore, might provide deeper insights into the underlying molecular mechanisms that regulate tumor microenvironment and tumorigenesis.

## DISCUSSION

Here, we introduced BayesDeep, a Bayesian hierarchical model for the reconstruction of gene expression at the single-cell resolution for NGS-based SRT data. BayesDeep links the cellular morphological information extracted from histology images and the spot-resolution molecular profiles derived from SRT data, enabling the inference of single-cell-resolution molecular profiles. Specifically, BayesDeep employs a negative binomial distribution to model spot-resolution gene expression and considers the latent normalized gene expression as an average of single-cell-resolution gene expression across all cells within a given spot. Our findings demonstrated the superior performance of BayesDeep compared to other competing methods for gene expression reconstruction in both simulation study and real SRT data.

Furthermore, by applying BayesDeep to two real SRT data, we unveiled detailed expression patterns that cannot be captured at the spot resolution, which offers invaluable insights for further exploration of the subsequent biomedical research. Firstly, cell populations can be separated to characterize different cell types or subpopulations based on the predicted single-cell-resolution molecular profiles, which can be further applied to marker gene identification and automated annotation^40^. Secondly, pseudotime analysis on the reconstructed high-resolution molecular profiles helps to understand the temporal progression of cellular states or trajectories within the tumor microenvironment, which further aids in unraveling the dynamic nature of tumors, providing valuable information on how cells change and evolve over time^41^. Last but not least, reconstructed molecular profiles at the single-cell resolution enable the examination of cell-cell communication patterns, which further contributes to a deeper understanding of cellular behavior, intercellular signaling dynamics, and their implications in clinical studies^32^. In summary, the downstream analysis indicates the significance of BayesDeep in exploring the tumor microenvironment and advancing the study of molecular biology.

There are several important future extensions for BayesDeep. First, the performance of BayesDeep highly relies on the selection of cellular features included in the model. On the simulated data where the actual gene expression is artificially generated and thus totally determined by the included covariates, BayesDeep can reconstruct single-cell-resolution gene expressions highly consistent with the actual expression. However, for the model validation on mouse visual cortex STARmap data, the limited cellular characteristic data available may constrain a comprehensive reconstruction of gene expression, thus resulting in relatively low consistency with the actual gene expression. Therefore, improving nuclei segmentation and classification methods offers more accurate cell biological and morphological characteristics and, in turn, further promotes the performance of BayesDeep. Second, it is necessary to optimize the computational efficiency of BayesDeep for large-scale studies, especially for larger data that are becoming increasingly common with the advancement of SRT technologies. In addition, there is potential for the BayesDeep model to integrate external data or other types of omics data, such as scRNA-seq data, to enhance gene expression inference. Integrating other data types could provide a more comprehensive understanding of the cellular environment and the complex interactions within cells. Lastly, we anticipate that BayesDeep could be extended to generate molecular profiles for three-dimensional tissue regions. We have demonstrated BayesDeep’s ability to extrapolate gene expression in a region with a similar cellular environment but without SRT data. With the available three-dimensional image profiles, BayesDeep can be adjusted to build molecular profiles at the single-cell resolution on three-dimensional tissue sections and provide more detail and context to cellular interactions and tissue architecture. These future directions could further boost the performance and generalizability of BayesDeep.

## METHODS

In this section, we define the spot-resolution molecular profile of NGS-based SRT data generated from ST or the improved 10x Visium platform and the single-cell-resolution image profile derived from the AI-reconstructed histology image paired with the SRT data. Then, we describe the geospatial profile that captures the spatial relationship between spots and cells. Following this, we detail the Bayesian statistical model, BayesDeep, employed for the reconstruction of expression for each gene of interest at the single-cell resolution. For quick reference, all data and parameter notations introduced in this paper are summarized in Supplementary Table S2.

### Data Preparation

The spot-resolution molecular profile. We denote the SRT molecular profile (i.e., gene expression measurements in terms of counts) as an *N*-by-*P* matrix ***Y*** = J*y*_*ij*_]_*N*×*P*_, where each entry *y*_*ij*_ ∈ ℕ represents the read count for gene *j* (*j* = 1, •, *P*, with *P* being the total number of genes) observed at spot *i* (*i* = 1, •, *N*, with *N* being the total number of spots). These *N* spots are regularly arrayed on a two-dimensional square or triangle lattice, with their spatial coordinates given by an *N*-by-2 matrix 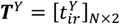, where each row 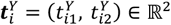 records the x and y-coordinates of spot *i* in the designated domain.

#### The single-cell-resolution image profile

We denote the image profile, derived from the histology image paired with the SRT data, as an *M*-by-*L* matrix ***X*** = [*x*_*ml*_]_*M*×*L*_, where each entry *x*_*ml*_ ∈ ℝ represents a measurement for explanatory variable *l* (*l* = 1, •, *L*, with *L* being the total number of explanatory variables) observed for cell *m* (*m* = 1, •, *M*, with *M* being the total number of cells). Our study utilizes *L* single-cell-resolution morphological features as the explanatory variables, extracted using the histology-based digital (HD)-staining model^13^. Specifically, the HD-staining model is a deep-learning model based on the mask regional convolutional neural network (Mask R-CNN) architecture^42^, which is trained to segment the nucleus of various cell types, such as immune, tumor, and stromal cells. HD-staining then computes ten geometric shape features for each identified cell nucleus, including filled area, net area, convex area, extent, perimeter, solidity, eccentricity, major axis length, minor axis length, and orientation. The definitions of these shape features can be found in Table S3 of a recent study^43^. Supposing there are *Q* different types of cells coded in *Q* dummy variables, the total number of explanatory variables is *L* = *Q* + 10. The HD-staining model was initially trained using histology images from lung adenocarcinoma patients in the National Lung Screening Trial (NLST) study (https://biometry.nci.nih.gov/cdas/nlst/), wherein nuclei of six different cell types were manually annotated by well-experienced pathologists. Although the original model was trained by using data specific to lung cancer, it has been improved and validated to adapt to histology images from various cancer types, including breast cancer, prostate cancer, and other carcinomas. To represent cell nuclei locations, we use an *M*-by-2 matrix 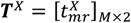, where each row 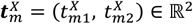 records the x and y-coordinates of cell *m* in the designated domain.

#### The spot-cell geospatial profile

In the context of NGS-based SRT technologies, spots are defined as circular regions comprising barcoded mRNA capture probes, where gene expression is quantified within a given tissue section^3^. The SRT data provides both the physical spot diameter *d* and its corresponding length in pixels. Assuming that the x and y-coordinates in spot-resolution geospatial profile *T*^*Y*^ represent the locations of spot centers, then we can identify whether cell *m* is located within spot *i* by a direct evaluation of the condition 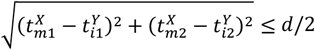. This allows us to construct the spot-cell geospatial profile, denoted by an *N*-by-*M* binary matrix ***G*** = [*g*_*im*_]_*N*×*M*_, which reflects whether a given cell *m* is within the barcoded area of a given spot *i* (i.e., *g*_*im*_ = 1) or not (i.e., *g*_*im*_ = 0). Notably, the coverage of measured areas in NGS-based SRT techniques is relatively limited, encompassing approximately 38% of the overall tissue section. As a result, most cells do not fall within the boundaries of any of the defined spots. In our real data analysis, for instance, 97,135 out of *M* = 156,115 cells for the human breast cancer 10x Visium data, and 252,765 out of *M* = 352,818 cells for the human prostate cancer 10x Visium data, are not covered by the measured area of any spots. This limitation in the scope of measured regions can lead to substantial information loss when analyzing SRT data. This motivates us to develop BayesDeep to address this challenge and enhance the analysis of SRT data.

### Model Description

BayesDeep is essentially a negative binomial regression model with regularization for handling grouped observations under the Bayesian framework. Its primary objective is to utilize the spot-level molecular profile, single-cell-resolution image profile, and spot-cell geospatial profile to reconstruct the single-cell-resolution molecular profile, which can be represented by an *M*-by-*P* matrix ***Θ*** = [θ_*mj*_]_*M×P*_, where each entry θ_*mj*_ ∈ ℝ^+^ is the predictive relative expression for gene *j* within cell *m*. It is worth noting that BayesDeep focuses on reconstructing relative gene expression at the single-cell resolution for one gene at a time (i.e., a specific column vector in ***Θ***), which allows for efficient parallel processing of multiple genes.

#### Modeling the spot-resolution molecular profile

We start by modeling the over-dispersed spot-resolution gene expression matrix ***Y*** using a negative binomial (NB) distribution. NB-based models are widely used for analyzing sequence count data^44-47^ due to their ability to accommodate inherent over-dispersion. Specifically, we postulate that each observed read count follows an NB distribution:

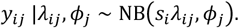

Here the NB(*v, Φ*) distribution is parameterized in terms of its mean *v* ∈ ℝ^+^ and dispersion 1/*Φ* ∈ ℝ^+^, with the probability mass function given by 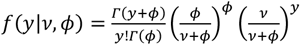. This parameterization provides the flexibility to characterize an unknown mean-variance structure, with the variance calculated *v* + *v*^*^/*Φ*. A small value of *Φ* indicates a high variance to mean ratio (i.e., 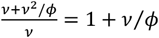, while a large value approaching infinity reduces the NB distribution to a Poisson distribution with the same mean and variance. The NB mean is further decomposed of two multiplicative components, 1) the size factor, denoted as *s*_*i*_, and 2) the spot-resolution relative expression for gene *j* observed at spot *i*, denoted as *λ*_*ij*_. Such a multiplicative characterization of the NB or Poisson mean is typical in both the frequentist^48-50^ and the Bayesian^51,52^ literature when modeling sequence count data. The set of *n* size factors is represented as ***s*** = [*s*_*i*_]_*N*×1_, capturing a range of biological and technical variabilities across samples, such as reverse transcription efficiency, amplification/dilution efficiency, and sequencing depth^53^. To ensure identifiability between these two parameters, we set *s*_*i*_ proportional to the summation of the total read counts across all genes at spot *i* ^54^. With a constraint of 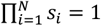, we compute 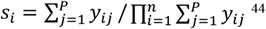^44^. It is noteworthy that alternative methods for estimating ***s*** are available, such as setting each *s*_*i*_ to be proportional to the upper quartiles of non-zero read counts across all genes at spot *i* ^55^, or even modeling ***s*** through a Dirichlet process mixture model with mean constraint^56^. To complete the prior model specification, we place a common gamma prior on all dispersion parameters, denoted as *Φ* = [*Φ*_*j*_]_*P*×1_, that is, *Φ*_*j*_ ∼ Ga(*a*_ϕ_, *b*_ϕ_), where *a*_ϕ_ and *b*_ϕ_ are fixed hyperparameters. We recommend choosing small values, such as *a*_ϕ_ = *b*_ϕ_ = 0.1, to maintain a weakly informative setting.

#### Linking the spot-resolution and single-cell-resolution molecular profiles

We posit that the spot-resolution relative expression for gene *j* (i.e., *λ*_*ij*_) can be derived as an average of single-cell-resolution relative expression (i.e., θ_*mj*_’s) across all cells within the given spot *i*. This relationship can be formally expressed as:

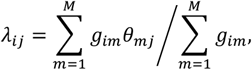

where the indicator variable *g*_*im*_ in the spot-cell geospatial profile ***G*** takes the value one if cell *m* is located within the barcoded area of spot *i*, and zero otherwise. The denominator in the above formulae, 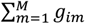, represents the total number of cells within spot *i* observed in the paired histology images. From an alternative perspective, this assumption can be interpreted as the spot-resolution gene expression being an average of cell-type-specific gene expression within the same spot, weighted by the respective cell-type proportions. Notably, this foundational assumption finds validations in various cell-type deconvolution algorithms designed for SRT data^57-59^. From a statistical viewpoint, this assumption enables the application of a regression model tailored to accommodate grouped observations^60,61^.

#### Linking the single-cell-resolution molecular and image profiles

In our pursuit of predicting the single-cell-resolution gene expression θ_*mj*_ ‘s, we leverage morphological information encompassing cell type and nuclei-shape features of each cell *m*. Specifically, we adopt a linear model to capture the relative expression for gene *j* within cell *m*:

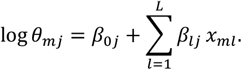

The choice of a log-link function is driven by the necessity for maintaining θ_*mj*_ values in the positive domain, thereby ensuring a positive NB mean. In this formulation, *β*_0*j*_ is the baseline expression for gene *j* shared by all cells. Note that exp(*β*_0*j*_) can also be interpreted as a scaling factor that adjusts for gene-specific effects. As previously introduced, we use the *M*-by-*L* matrix ***X*** to present observations from *L* explanatory variables extracted from the histology image, including cell type information, cell nuclei-shape descriptors, and other important single-cell-resolution measurements relevant to the analysis. Given the coding of *Q* cell types into *Q* dummy variables, we added a constraint 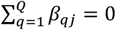 to avoid identifiability issues arising from the sum of the components. Consequently, exp(*β*_*lj*_), corresponding to cell type *Q*, represents the mean relative expression for gene *j* across all cells. Each column in the *L*-by-*P* coefficient matrix ***B*** = [*β*_*lj*_]_*L*×*P*_, denoted as ***β***_*j*_ = [*β*_*lj*_]_L×1_ with *β*_*lj*_ ∈ 󒄝, describes the effect of the *L* explanatory variables on the logarithm of the relative expression across cells for gene *j*. Hence, we can use the coefficient matrix ***B*** to explore the association between the image and molecular profiles at the single-cell resolution. In practice, for a specific gene *j*, it is likely that only a limited number of explanatory variables account for its expression. For instance, the expression of some housekeeping genes exhibits small variability among different tissues, cell types, or samples^62^. Under such circumstances, their corresponding coefficients *β*_*lj*_’s tend to be zeros. On the other hand, some pairs of explanatory variables might be highly correlated, leading to an inflation of regression coefficients and potentially harming prediction performance. We employ a regularization mechanism to prevent over-fitting and reduce the potential multicollinearity by specifying a spike-and-slab prior^63^ on each *β*_*lj*_. This prior is a mixture of distributions:

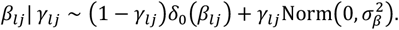

The spike component *δ*_0_(*β*_*lj*_) is a point mass distribution at *β*_*lj*_ = 0, while the slab component is a normal distribution centered at zero. If the auxiliary binary variable *γ*_*lj*_ = 1, then the probability of *β*_*lj*_ = 0 is zero, indicating that explanatory variable *l* is relevant for explaining the relative expression for gene *j*. Conversely, *γ*_*lj*_ = 0 restricts that *β*_*lj*_ = 0, indicating that explanatory variable *l* has no contribution to the gene *j*’s expression. This prior setting enables us to identify significant associations between gene expression and explanatory variables, which, in our case, are the morphological features. The variance of the slab component 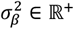 is a fixed hyperparameter set to one. We complete the model specification by placing an independent Bernoulli distribution on *γ*_*lj*_, i.e., *γ*_*lj*_ ∼ Bern(*π*_γ_), where *π*_γ_ ∈ (0,1) is a fixed hyperparameter that indicates the percentage of explanatory variables included in the final model *a priori*. We set *π*_γ_ = 0.5 to incorporate relatively weak information.

#### Full data likelihood and posterior

The model parameter space consists of 1) the dispersion parameter *Φ* that accounts for the over-dispersion commonly observed in gene expression data, 2) the coefficient matrix ***B*** that quantifies the relationship between gene expression as measured in SRT data and the morphological features extracted from the paired histology image, and 3) the *L*-by-*P* selection matrix ***Γ*** = [*γ*_*lj*_]_*L*×*P*_ that indicates the significant association in the coefficient matrix ***B***. The complete data likelihood can be written as:

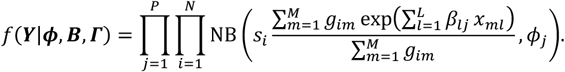

With the prior specifications detailed above, the full posterior distribution can be written as:

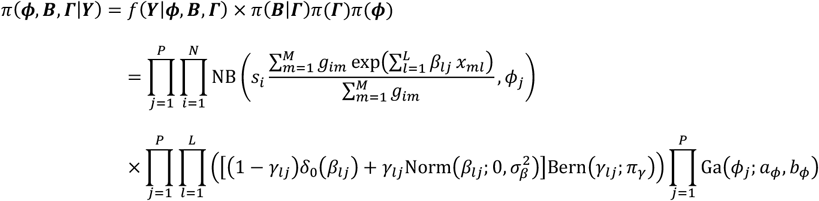

### Model Fitting

#### Posterior sampling *via* MCMC algorithms

We explore the posterior distribution *via* a Markov chain Monte Carlo (MCMC) algorithm based on stochastic search variable selection^64,65^. Specifically, we iteratively update each parameter using a Metropolis-Hasting (MH) algorithm. We note that this algorithm is sufficient to guarantee ergodicity for our model. Full details regarding the implementation of the MCMC algorithm are available in Section S2 in Supplementary Materials.

#### Posterior inference

We obtain posterior inference by post-processing the MCMC samples following the burn-in phase. Let 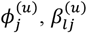 and 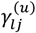 denote the posterior samples in the *u*-th iteration after burn-in, where *u* = 1, …, *U*. Our primary focus is on the selection of the important explanatory variables for each gene *j, via* the selection matrix *Γ*. An effective approach to summarize the posterior distributions of these binary parameters is by computing the marginal posterior probability of inclusion (PPI):

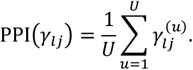

Then, we set 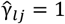 if its corresponding PPI exceeds a pre-specified threshold. When choosing the threshold, we recommend either opting for a threshold of 0.5, leading to a median model, or following a procedure that controls the expected Bayesian false discovery rate ^66^. For each dispersion parameter *Φ*_*j*_ and each coefficient *β*_*lj*_, we estimate them by calculating their posterior means by averaging over all their respective MCMC samples after burn-in,

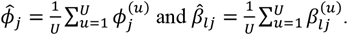

Additionally, a quantile estimation or credible interval for each parameter of interest can be obtained from MCMC samples.

#### Predictive inference

Our primary goal is to reconstruct gene expression at the single-cell resolution by estimating the matrix ***Θ*** = [θ_*mj*_]_*M*×*P*_. On the basis of the MCMC samples on the coefficient matrix ***B*** = [*β*_*lj*_]_*L*×*P*_, we predict each θ_*mj*_ by Monte Carlo simulation. Specifically, at each iteration *u* after burn-in, we compute the relative gene expression for gene *j* within cell *m* as follows:

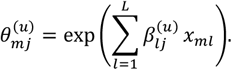

Subsequently, we sample the gene expression for gene *j* within spot *i* using an NB distribution,

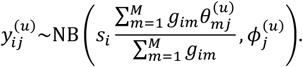

Consequently, both single-cell-resolution relative gene expression 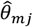 and spot-resolution gene expression 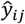 can be estimated by summarizing their corresponding MCMC samples. For instance, their predictive means can be approximated as follows:

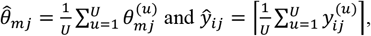

where ⌈·⌉ denotes the ceiling function.

## Data availability

The authors analyzed two publicly available SRT data. Raw count matrices, images, and spatial information for two SRT data from 10x Visium are accessible on the 10x Genomics website at https://support.10xgenomics.com/spatial-gene-expression/datasets.

## Code availability

An open-source implementation of the BayesDeep algorithm in R/C++ is available at https://github.com/Xijiang1997/BayesDeep.

## Acknowledgments

This work was supported by the following funding: the National Science Foundation [2210912, 2113674] and the National Institutes of Health [1R01GM141519] (to Q. L.); the National Institutes of Health [R01GM140012, R01GM141519, R01DE030656, U01CA249245], and the Cancer Prevention and Research Institute of Texas [CPRIT RP180805, CPRIT RP230330] (to G. X.); the Rally Foundation, Children’s Cancer Fund (Dallas), the Cancer Prevention and Research Institute of Texas (RP180319, RP200103, RP220032, RP170152 and RP180805), and the National Institutes of Health (R01DK127037, R01CA263079, R21CA259771, UM1HG011996, and R01HL144969) (to L. X.). The funding bodies had no role in the design, collection, analysis, or interpretation of data in this study.

