## Supplementary materials for "Reconstructing Spatial Transcriptomics at the Single-cell Resolution with BayesDeep"

### Supplementary Materials for
### "Reconstructing spatial transcriptomics at the single-cell resolution with BayesDeep"

**Xi Jiang**[1,2], **Lei Dong**[1], **Shidan Wang**[1], **Zhuoyu Wen**[1], **Mingyi Chen**[3], **Lin Xu**[1,4,*], **Guanghua Xiao**[1,*], and **Qiwei Li**[5,*]

[1]Quantitative Biomedical Research Center,
Peter O'Donnell Jr. School of Public Health,
The University of Texas Southwestern Medical Center,
Dallas, TX 75390, United States.

[2]Department of Statistics and Data Science,
Southern Methodist University,
Dallas, TX 75275, United States.

[3]Department of Pathology,
The University of Texas Southwestern Medical Center,
Dallas, TX 75390, United States.

[4]Department of Pediatrics, Division of Hematology/Oncology,
The University of Texas Southwestern Medical Center,
Dallas, TX 75390, United States.

[5]Department of Mathematical Sciences,
The University of Texas at Dallas,
Richardson, TX 75080, United States.

#### S1. Simulated Data Generative Model

In our simulation study, we aimed to validate the accuracy of our model. We selected $N = 500$ adjacent spots from the human breast cancer 10x Visium data (see Figure 2A in the manuscript) with known spot locations $\boldsymbol{T}^Y$. The cell locations $\boldsymbol{T}^X$ and the covariate matrix $\boldsymbol{X}$, including cell types and nuclei shape features, were obtained from the nuclei identification results through HD-staining on the paired histology image. As detailed in Methods Section of the manuscript, in real SRT data, typically only a subset of covariates correlate with gene expression for most genes. To replicate this scenario, we generated coefficients with high sparsity. Specifically, for cell-type-related explanatory variables, we assumed all corresponding coefficients to be non-zero and normally distributed as $\beta_{lj} \sim \mathrm{N}(1, 1)$. In contrast, for nuclei-shape-related explanatory variables, we produced highly sparse coefficients as follows:

$$\beta_{lj} \sim 0.9I(\beta_{lj} = 0) + 0.1[0.5\mathrm{Unif}(-1, -0.5) + 0.5\mathrm{Unif}(0.5, 1)].$$

Consequently, the coefficient matrix $\boldsymbol{B}$ exhibited 90% sparsity. We then generated single-cell-resolution expression levels in line with our model assumption $\boldsymbol{\theta}_j = \exp(\boldsymbol{X}\boldsymbol{\beta}_j)$. Spot-resolution relative expression levels for gene $j$ (i.e., $\boldsymbol{\lambda}_j$) were calculated as $\boldsymbol{\lambda}_j = (\boldsymbol{G}\boldsymbol{\theta}_j)/(\boldsymbol{G}\mathbf{1})$, where $\boldsymbol{G}$ is derived from spot and cell locations (i.e., $\boldsymbol{T}^Y$ and $\boldsymbol{T}^X$), and the denominator $\boldsymbol{G}\mathbf{1}$ represents the number of cells covered by each spot. To simulate observed spot-resolution gene expression counts, we followed the process outlined in Sun et al. (2020) and Jiang et al. (2022). Specifically, we generated the size factor $s_i \sim \mathrm{Unif}(0.5, 1.5)$ for $i = 1, \ldots, N$ and dispersion parameter $\phi_j \sim \mathrm{Exp}(0.1)$ for $j = 1, \ldots, P$, and subsequently, each read count $y_{ij} \sim \mathrm{NB}(s_i\lambda_{ij}, \phi_j)$. This data generation process was repeated for $P = 300$ genes. Finally, with the generated $\boldsymbol{Y}$ and the observed $\boldsymbol{X}$ and $\boldsymbol{G}$, we examined our model's ability in reconstructing the single-cell-resolution molecular profile $\boldsymbol{\Theta}$.

#### S2. Details of the MCMC Algorithms

The model parameter space $(\boldsymbol{\phi}, \boldsymbol{B}, \boldsymbol{\Gamma})$ consists of 1) the NB dispersion parameters $\boldsymbol{\phi}$ that accounts for the over-dispersion commonly observed in gene expression count data, 2) the coefficient matrix $\boldsymbol{B}$ that quantifies the relationship between gene expression as measured in SRT data and the morphological features extracted from the paired histology image, and 3) the selection matrix $\boldsymbol{\Gamma}$ that indicates the significant association in the coefficient matrix $\boldsymbol{B}$. The full likelihood of the model is as follows:

$$\begin{aligned} f(\boldsymbol{Y}|\boldsymbol{\phi}, \boldsymbol{B}, \boldsymbol{\Gamma}) &= \prod_{j=1}^{P} f(\boldsymbol{y}_j|\phi_j, \boldsymbol{\beta}_j, \boldsymbol{\gamma}_j) \\ &= \prod_{j=1}^{P}\prod_{i=1}^{N} f(y_{ij}|\phi_j, \boldsymbol{\beta}_j) \\ &= \prod_{j=1}^{P}\prod_{i=1}^{N} \mathrm{NB}\left(y_{ij}; s_i \frac{\sum_{m=1}^{M} g_{im} \exp\left(\sum_{l=1}^{L} \beta_{lj} x_{ml}\right)}{\sum_{m=1}^{M} g_{im}}, \phi_j\right). \end{aligned} \tag{1}$$

With the prior specifications detailed in Methods Section of the manuscript, we give the full posterior distribution as follows:

$$\begin{aligned} \pi(\boldsymbol{\phi}, \boldsymbol{B}, \boldsymbol{\Gamma}|\boldsymbol{Y}) &= f(\boldsymbol{Y}|\boldsymbol{\phi}, \boldsymbol{B}, \boldsymbol{\Gamma}) \times \pi(\boldsymbol{B}|\boldsymbol{\Gamma})\pi(\boldsymbol{\Gamma})\pi(\boldsymbol{\phi}) \\ &= f(\boldsymbol{Y}|\boldsymbol{\phi}, \boldsymbol{B}, \boldsymbol{\Gamma}) \times \left(\prod_{j=1}^{P}\prod_{l=1}^{L} \pi(\beta_{lj}|\gamma_{lj})\right)\left(\prod_{j=1}^{P}\prod_{l=1}^{L} \pi(\gamma_{lj})\right)\left(\prod_{j=1}^{P} \pi(\phi_j)\right) \\ &= \prod_{j=1}^{P}\prod_{i=1}^{N} \mathrm{NB}\left(y_{ij}; s_i \frac{\sum_{m=1}^{M} g_{im} \exp\left(\sum_{l=1}^{L} \beta_{lj} x_{ml}\right)}{\sum_{m=1}^{M} g_{im}}, \phi_j\right) \\ &\times \prod_{j=1}^{P}\prod_{l=1}^{L} (1-\gamma_{lj})\delta_0(\beta_{lj}) + \gamma_{lj}\mathrm{N}(\beta_{lj}; 0, \sigma_\beta^2) \\ &\times \prod_{j=1}^{P}\prod_{l=1}^{L} \mathrm{Bern}(\gamma_{lj}; \pi_\gamma) \\ &\times \prod_{j=1}^{P} \mathrm{Ga}(\phi_j; a_\phi, b_\phi) \end{aligned} \tag{2}$$

MCMC algorithm was conducted for the posterior sampling. Specifically, we performed the following steps sequentially at each MCMC iteration after a random initialization.

**Update of over-dispersion parameter $\boldsymbol{\phi}$**: We updated each $\phi_j$ for $j = 1, \ldots, P$, independently by using the random walk Metropolis-Hasting algorithm. We first proposed a new $\phi_j^*$ from the log-normal distribution, i.e., $\ln(\phi_j^*) \sim \mathrm{N}(\ln(\phi_j), \tau_\phi^2)$, and accepted the proposed value with probability $\min(1, r_\phi)$, where

$$r_\phi = \frac{\prod_{i=1}^N f(y_{ij}|\phi_j^*, \boldsymbol{\beta}_j) \times \pi(\phi_j^*) \times J(\phi_j; \phi_j^*)}{\prod_{i=1}^N f(y_{ij}|\phi_j, \boldsymbol{\beta}_j) \times \pi(\phi_j) \times J(\phi_j^*; \phi_j)}.$$

Note that the proposal density ratio equals to 1 for this random walk Metropolis update on $\phi_j$.

**Joint update of covariate coefficient $\boldsymbol{B}$ and the selection indicator $\boldsymbol{\Gamma}$**: We updated each $\beta_{lj}$ and $\gamma_{lj}$ for $l = 1, \ldots, L$ and $j = 1, \ldots, P$, sequentially by using an independent Metropolis-Hasting algorithm. A between-model step was implemented first to jointly update $\beta_{lj}$ and $\gamma_{lj}$. We used the *add-delete* algorithm, where we changed the binary value of $\gamma_{lj}$ to $1 - \gamma_{lj}$ each time. We repeated this step multiple times to ensure validation of the feature selection.

For the add case, i.e., changing from $\gamma_{lj} = 0$ to $\gamma_{lj}^* = 1$, we proposed $\beta_{lj}^*$ from the normal distribution $\mathrm{N}(0, \tau_\beta^2)$. For the delete case, i.e., $\gamma_{lj} = 1 \ \rightarrow \ \gamma_{lj}^* = 0$, we set $\beta_{lj}^* = 0$. We finally accepted the proposed $\gamma_{lj}^*$ with probability $\min(1, r_\gamma)$, where

$$r_\gamma = \frac{\prod_{i=1}^N f(y_{ij}|\phi_j,\beta_{lj}^*,\boldsymbol{\beta}_{-l,j})\times\pi(\beta_{lj}^*|\gamma_{lj}^*)\times\pi(\gamma_{lj}^*)\times J(\beta_{lj};\beta_{lj}^*|\gamma_{lj};\gamma_{lj}^*)\times J(\gamma_{lj};\gamma_{lj}^*)}{\prod_{i=1}^N f(y_{ij}|\phi_j,\beta_{lj},\boldsymbol{\beta}_{-l,j})\times\pi(\beta_{lj}|\gamma_{lj})\times\pi(\gamma_{lj})\times J(\beta_{lj}^*;\beta_{lj}|\gamma_{lj}^*;\gamma_{lj})\times J(\gamma_{lj}^*;\gamma_{lj})},$$

where the last proposal density ratio equals to 1 and the second to last proposal density ratio equals to $1/N(\beta_{lj}^*; 0, \tau_\beta^2)$ for the add step and $N(\beta_{lj}; 0, \tau_\beta^2)$ for the delete step.

**Update of covariate coefficient $\boldsymbol{B}$**: We performed a within-model step to update each $\beta_{lj}$ that corresponds to $\gamma_{lj} = 1$ to improve MCMC mixing. We first proposed a new $\beta_{lj}^*$ from the normal distribution $\mathrm{N}(\beta_{lj}, (\sigma_\beta/2)^2)$ and then accepted the proposed value with probability $\min(1, r_\beta)$, where

$$r_\beta = \frac{\prod_{i=1}^N f(y_{ij}|\phi_j,\beta_{lj}^*,\boldsymbol{\beta}_{-l,j})\times\pi(\beta_{lj}^*|\gamma_{lj}{=}1)\times J(\beta_{lj};\beta_{lj}^*)}{\prod_{i=1}^N f(y_{ij}|\phi_j,\beta_{lj},\boldsymbol{\beta}_{-l,j})\times\pi(\beta_{lj}|\gamma_{lj}{=}1)\times J(\beta_{lj}^*;\beta_{lj})}.$$

Note that the proposal density ratio equals to 1 for this random walk Metropolis update on $\beta_{lj}$.

#### S3. Implementation Details of Competing Methods

We adapted the TESLA workflow described in their online tutorial (`https://github.com/jianhuupenn/TESLA/blob/main/tutorial/tutorial.md`). Specifically, we normalized the gene expression measurements into log scale. For the histology image, we detected tissue contour using *cv2*. All parameters were kept at the defaults or as the same as in the tutorial. We then imputed the gene expression on all pixels within the contour. The predicted gene expression at each cell was set to be the imputed gene expression on the pixel where the cell is located.

Gaussian process (GP) was implemented with *GaussianProcessRegressor* from *sklearn.gaussian_process* module in *Python*. Before fitting the GP model, the observed read counts $\mathbf{y}_j$ for gene $j$ were normalized to $\tilde{y}_j = \log(\mathbf{y}_j/\mathbf{s} + 1)$, where $\mathbf{s} = (s_1, \ldots, s_N)$ was the estimated size factor of spots and was computed as $s_i \propto \sum_{j=1}^{P} y_{ij}$. The covariance function of the GP was specified using the radial basis function (RBF) kernel, with the length scale of the kernel set to be the radius of spots. Additionally, to account for noise, a *WhiteKernel* component was incorporated into the kernel, allowing for the estimation of the global noise level from the data. The GP model was then trained utilizing the location information at spot resolution as features and the normalized gene expression as response variable. For predicting gene expression at the single-cell resolution, we input the location information of all cells and employed the mean of the output predictive distribution as the predicted gene expression value.

#### REFERENCES

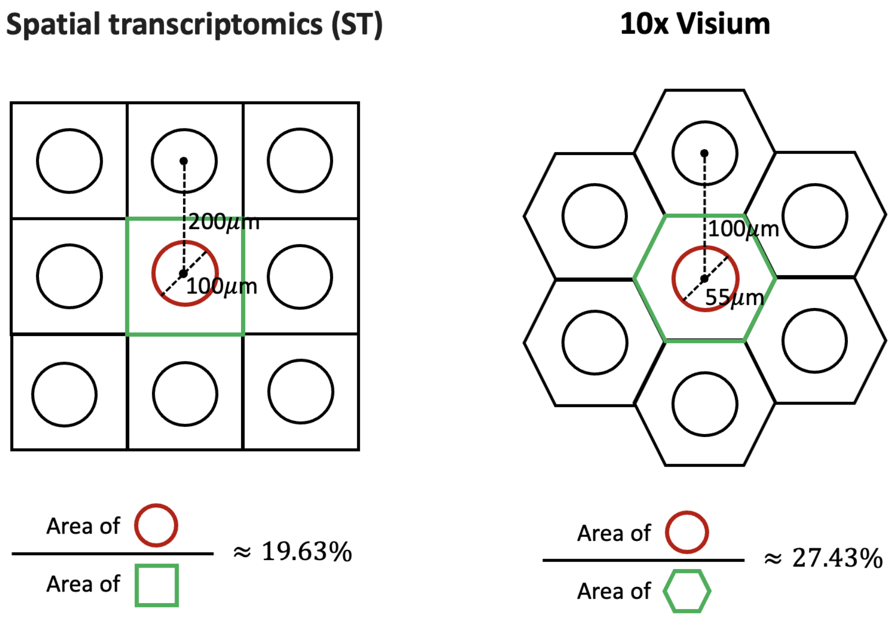


**Figure S1**: Geometric representations of a barcoded spot (in red) and its expanded region (in green) on the spatial transcriptomics (ST) and the improved 10x Visium platforms. For ST, the spot diameter is $55\mu m$ with a center-to-center distance of $100\mu m$ between two adjacent spots. For 10x Visium, these measures are $100\mu m$ and $200\mu m$, respectively. The percentage of area covered by ST barcoded spots can be approximated by the ratio of the red circle area to the green square, calculated as $\frac{\pi\times\left(\frac{100\mu m}{2}\right)^2}{(200\mu m)^2} = 19.63\%$. Similarly, the percentage of area covered by 10x Visium barcoded spots can be approximated by the ratio of the red circle to the green hexagon, calculated as $\frac{\pi\times\left(\frac{55\mu m}{2}\right)^2}{\frac{3\sqrt{3}}{2}\times\left(\frac{2}{\sqrt{3}}\frac{100\mu m}{2}\right)^2} = 27.43\%$.

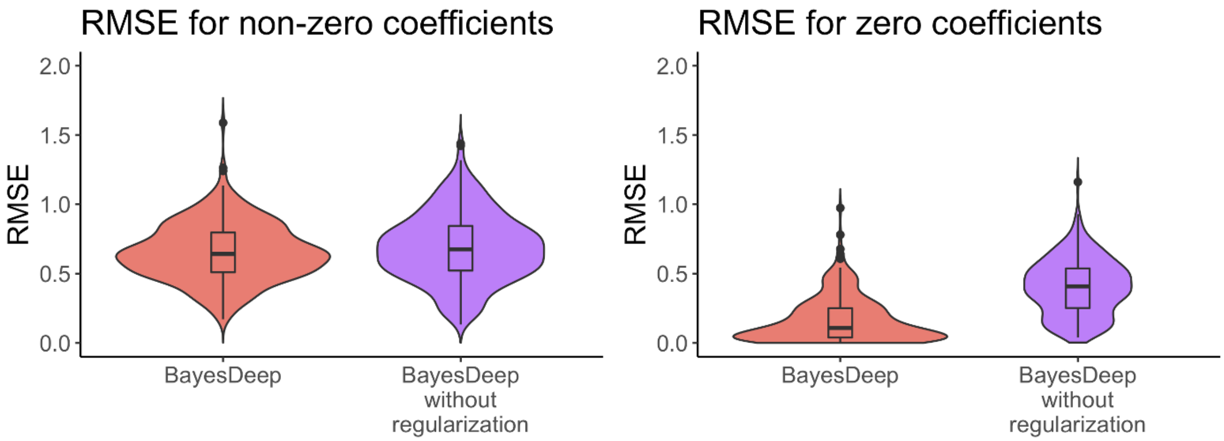


**Figure S2**: Overview of model validation for the simulated data: Evaluation results in terms of the root mean square error (RMSE) between the actual and estimated covariate coefficient $\beta_{lj}$'s for BayesDeep with and without regularization, respectively.

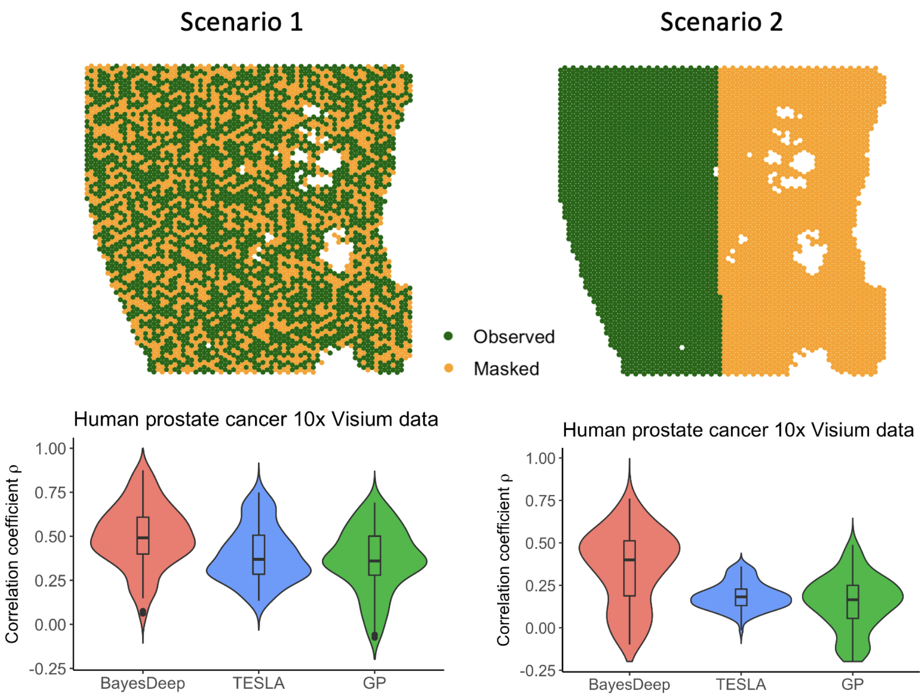


**Figure S3**: Overview of model validation for the human prostate cancer 10x Visium data, including the validation settings and evaluation results in terms of the Pearson correlation coefficients $\rho$ between the actual and predicted gene expression counts ($y_{ij}$'s vs. $\hat{y}_{ij}$'s) for BayesDeep, TESLA, and Gaussian Process (GP), respectively.

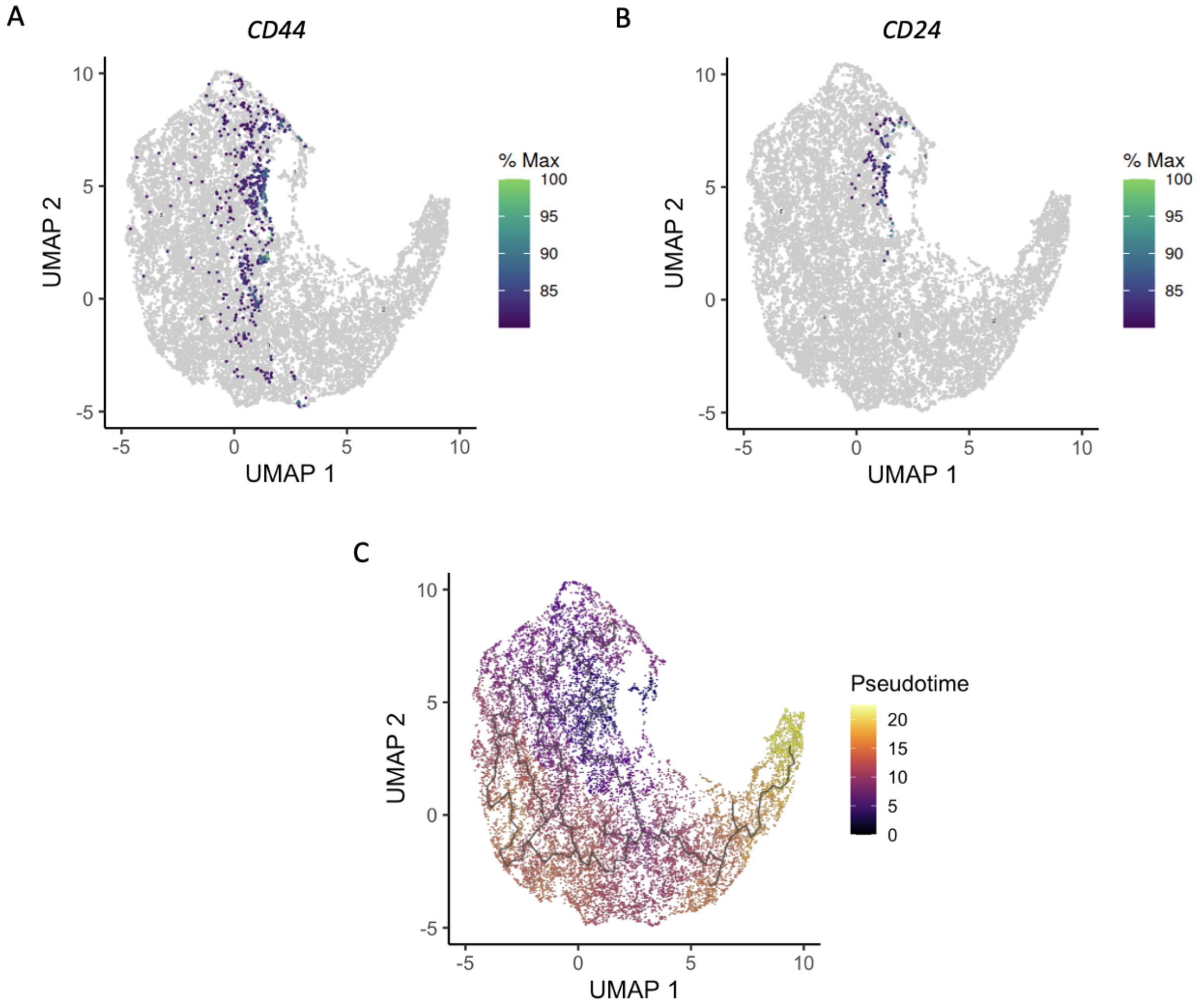


**Figure S4**: Downstream analysis on the human breast cancer 10x Visium data: A. BayesDeep-generated single-cell-resolution expression of the breast cancer stem cell markers *CD44* depicted on the UMAP. B. BayesDeep-generated single-cell-resolution expression of the breast cancer stem cell markers *CD24* depicted on the UMAP. C. Pseudotime analysis on the UMAP.

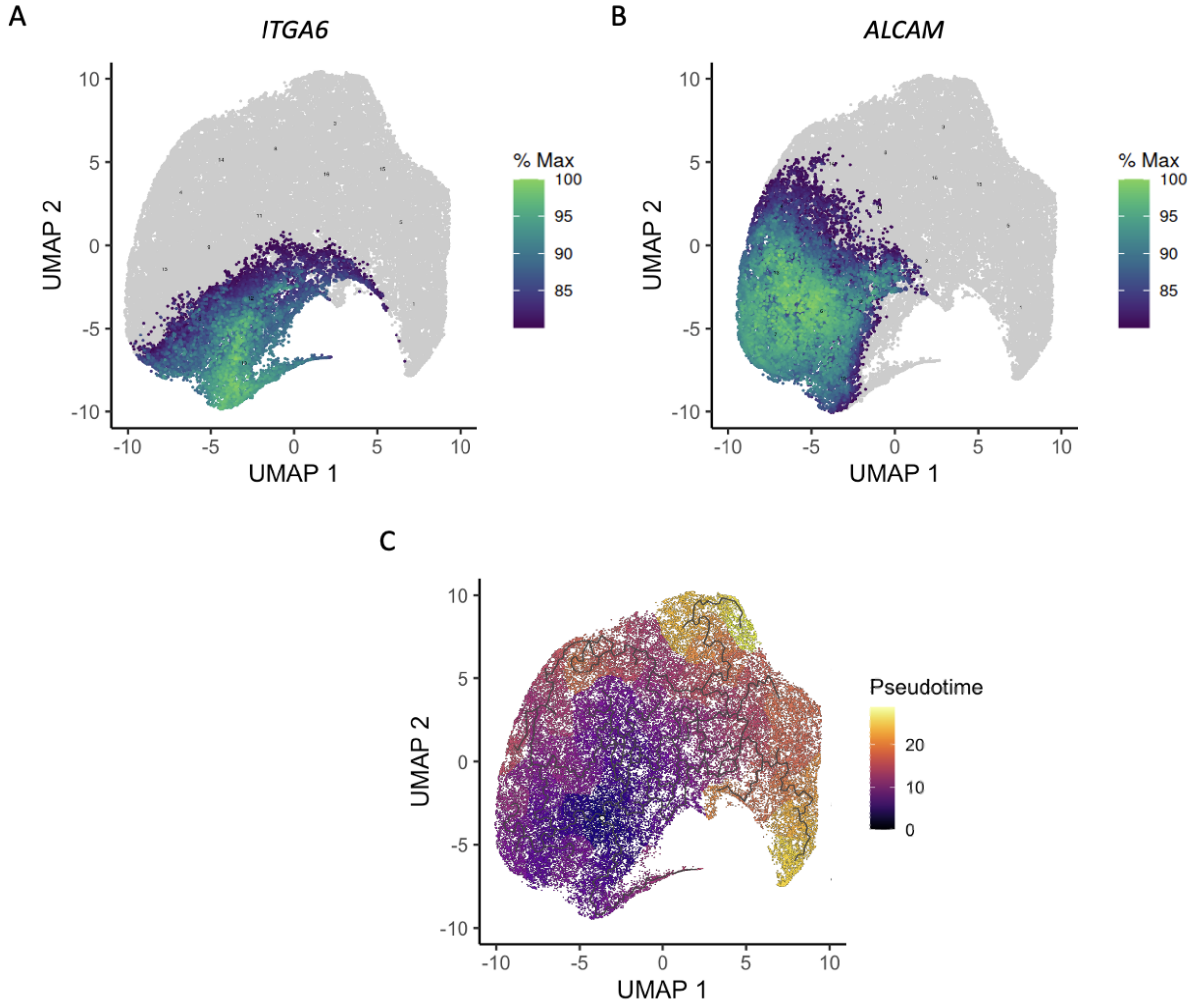


**Figure S5**: Downstream analysis on the human prostate cancer 10x Visium data: A. BayesDeep-generated single-cell-resolution expression of the prostate cancer stem cell markers *ITGA6* depicted on the UMAP. B. BayesDeep-generated single-cell-resolution expression of the prostate cancer stem cell markers *ALCAM* depicted on the UMAP. C. Pseudotime analysis on the UMAP.

Table S1: A list of explanatory variables utilized by BayesDeep in real data analysis, including cell type classification and nuclei shape features extracted from the paired histology image by HD-staining

| | Human breast cancer 10x Visium data | Human prostate cancer 10x Visium data |
|---|---|---|
| Cell type | Ductal epithelium<br>Lymphocyte<br>Macrophage<br>Necrosis<br>Red blood cell<br>Stromal cell<br>Tumor cell | Lymphocyte<br>Macrophage<br>Necrosis<br>Red blood cell<br>Stromal cell<br>Tumor cell |
| Nuclei shape feature | Area<br>Convex area<br>Eccentricity<br>Extent<br>Filled area<br>Major axis length<br>Minor axis length<br>Orientation<br>Perimeter<br>Solidity | Area<br>Convex area<br>Eccentricity<br>Extent<br>Filled area<br>Major axis length<br>Minor axis length<br>Orientation<br>Perimeter<br>Solidity |

Table S2: A list of key notations of the proposed BayesDeep.

| | Notation | Support | Definition |
|---|---|---|---|
| Data | $N$ | $\mathbb{N}$ | The number of spots |
| | $P$ | $\mathbb{N}$ | The number of genes |
| | $M$ | $\mathbb{N}$ | The number of cells |
| | $L$ | $\mathbb{N}$ | The number of explanatory variables |
| | $\boldsymbol{Y} = [y_{ij}]_{N\times P}$ | $y_{ij} \in \mathbb{N}$ | The gene expression measurement table with ench entry $y_{ij}$ being the read count for gene $j$ observed at spot $i$ |
| | $\boldsymbol{s} = [s_i]_{N\times 1}$ | $s_i \in \mathbb{R}^+$ | The spot-specific size factors |
| | $\boldsymbol{T}^Y = [t^Y_{ir}]_{N\times 2}$ | $(t^Y_{i1}, t^Y_{i2}) \in \mathbb{R}^2$ | The spot location with each row $(t^Y_{i1}, t^Y_{i2})$ being the $x$ and $y$ coordinates of spot $i$ |
| | $\boldsymbol{X} = [x_{ml}]_{M\times L}$ | $x_{ml} \in \mathbb{R}$ | The covariate matrix with each entry $x_{ml}$ representing a measurement for explanatory variable $l$ observed for cell $m$ |
| | $\boldsymbol{T}^X = [t^X_{mr}]_{M\times 2}$ | $(t^X_{m1}, t^X_{m2}) \in \mathbb{R}^2$ | The cell location with each row $(t^X_{i1}, t^X_{i2})$ being the $x$ and $y$ coordinates of cell $m$ |
| | $\boldsymbol{G} = [g_{im}]_{N\times M}$ | $g_{im} \in \{0,1\}$ | The spot-cell spatial relationship matrix with $g_{im} = 1$ indicating that cell $m$ locates within spot $i$ and $g_{im} = 0$ otherwise |
| Parameters | $\boldsymbol{\phi} = [\phi_j]_{P\times 1}$ | $\phi_j \in \mathbb{R}^+$ | The gene-specific negative binomial dispersion parameter |
| | $\boldsymbol{\Lambda} = [\lambda_{ij}]_{N\times P}$ | $\lambda_{ij} \in \mathbb{R}^+$ | The relative gene expression at the spot resolution with each entry $\lambda_{ij}$ being the relative gene expression for gene $j$ at spot $i$ |
| | $\boldsymbol{\Theta} = [\theta_{mj}]_{M\times P}$ | $\theta_{mj} \in \mathbb{R}^+$ | The relative gene expression at the single-cell resolution with each entry $\theta_{mj}$ being the relative gene expression for gene $j$ at cell $m$ |
| | $\boldsymbol{B} = [\beta_{lj}]_{L\times P}$ | $\beta_{lj} \in \mathbb{R}$ | The covariate coefficient matrix with each entry $\beta_{lj}$ signifying the association between explanatory variable $l$ and gene $j$ |
| | $\boldsymbol{\Gamma} = [\gamma_{lj}]_{L\times P}$ | $\gamma_{lj} \in \{0,1\}$ | The selection matrix with each entry $\gamma_{lj} = 1$ indicating the corresponding covariate coefficient $\beta_{lj} \neq 0$ while $\gamma_{lj} = 0$ indicating $\beta_{lj} = 0$ |
| Hyper-parameters | $a_\phi$ | $\{0\} \cup \mathbb{R}^+$ | The shape parameter in the gamma prior for each $\phi_j$ |
| | $b_\phi$ | $\{0\} \cup \mathbb{R}^+$ | The rate parameter in the gamma prior for each $\phi_j$ |
| | $\sigma^2_\beta$ | $\mathbb{R}^+$ | The variance parameter of the slab component in the spike-and-slab prior for each $\beta_{lj}$ |
| | $\pi_\gamma$ | $(0,1)$ | The parameter in the Bernoulli prior for each $\gamma_{lj}$ |
